# Macrophages modulate fibrosis during newt lens regeneration

**DOI:** 10.1101/2023.06.04.543633

**Authors:** Georgios Tsissios, Anthony Sallese, J. Raul Perez-Estrada, Jared A. Tangeman, Weihao Chen, Byran Smucker, Sophia C. Ratvasky, Erika Grajales-Esquivel, Arielle Martinez, Kimberly J. Visser, Alberto Joven Araus, Hui Wang, Andras Simon, Maximina H. Yun, Katia Del Rio-Tsonis

## Abstract

Previous studies indicated that macrophages play a role during lens regeneration in newts, but their function has not been tested experimentally. Here we generated a transgenic newt reporter line in which macrophages can be visualized *in vivo*. Using this new tool, we analyzed the location of macrophages during lens regeneration. We uncovered early gene expression changes using bulk RNAseq in two newt species, *Notophthalmus viridescens* and *Pleurodeles waltl*. Next, we used clodronate liposomes to deplete macrophages, which inhibited lens regeneration in both newt species. Macrophage depletion induced the formation of scar-like tissue, an increased and sustained inflammatory response, an early decrease in iris pigment epithelial cell (iPEC) proliferation and a late increase in apoptosis. Some of these phenotypes persisted for at least 100 days and could be rescued by exogenous FGF2. Re-injury alleviated the effects of macrophage depletion and re-started the regeneration process. Together, our findings highlight the importance of macrophages in facilitating a pro-regenerative environment in the newt eye, helping to resolve fibrosis, modulating the overall inflammatory landscape and maintaining the proper balance of early proliferation and late apoptosis.

## Introduction

A popular hypothesis, fueled by the observations that regeneration is gradually lost through ontogeny and evolution, suggests that adult mammals have developed a more robust adaptive immune response at the expense of regenerative capabilities [1]. However, NOD/SCID mice that exhibited T cell deficiency but maintained macrophage numbers failed to regenerate their hearts at neonatal stages and demonstrated signs of severe fibrosis, suggesting that modulating adaptive immunity alone is not sufficient for successful regeneration [2]. Furthermore, the theory of inverse relationship between regeneration and immune proficiency fails to explain why several mammals are capable of epimorphic-like regeneration during adulthood [3–5]. An alternative theory suggests that the interaction between immune cells and the local microenvironment influences the regeneration outcome, rather than the absence or presence of special immune and regenerative cells [6].

Macrophage involvement during tissue regeneration has been a subject of intense discussion in recent years [7–10]. There is now increasing evidence that macrophages play a necessary role during limb, fin, tail, spinal cord, and heart regeneration in a variety of species [11–20]. The regenerative process in the aforementioned tissues and organs is complex, requires the integration of multiple cell populations, or is not easily accessible for visualization [21–24]. This added level of complexity makes it difficult to extrapolate immune mechanisms in a regenerative context. On the other hand, the case of lens regeneration in newts involves the transdifferentiation of a single cell type, the iris pigment epithelial cell (iPEC) into lens cells [25–30]. The elegant simplicity of this process serves as a great platform to uncover the mechanism by which macrophages promote or interfere with scar-free healing and regeneration.

In the past, studies utilizing electron microscopy technology recognized that macrophages, which were first called “special amoeboid cells” due to their morphology, migrated inside the iris epithelium at 3 days post-lentectomy (dpl) and phagocytosed melanosomes that were discharged from iPECs during the dedifferentiation process [31–34]. In the present study, we developed and characterized a new *Pleurodeles waltl* transgenic line with eGFP-labeled macrophages to enable the study of macrophages during the regeneration process. Furthermore, we characterized the immune landscape of the iris after injury and investigated the role of macrophages in directing the early and late stages of lens regeneration. We demonstrated that macrophage depletion modulates the wound healing response and affects the regenerative outcome. In addition, we demonstrated that macrophages returning into the deleterious wound lesion were unable to resolve the inflammatory and fibrotic environment. However, a secondary injury alone or the addition of fibroblast growth factor 2 (FGF2) restarted scar resolution and the regeneration processes.

## Results

### mpeg1:eGFP^MHY-SIMON^: A new transgenic newt reporter line for macrophages

Macrophage expressed gene 1 (*mpeg1*) is expressed in macrophages and has been successfully employed for driving fluorescent reporters to label macrophages in several species [35, 36]. Seeking to employ a similar approach for labeling macrophages in *Pleurodeles waltl*, we created a transgenic line in which the orthologous *mpeg1* promoter drives the expression of an eGFP-encoding sequence. We selected the positive offspring of the founder animals to obtain our first generation (F_1_) of the *Pleurodeles waltl* transgenic line tgTol2(Dre.*mpeg1:eGFP*)^MHY/SIMON^ (referred as mpeg1:GFP from now on). To test the stability of our transgenic line, we screened all the mpeg1:GFP embryos from generation F_1_ onwards for fluorescent signals. In the positive offspring, eGFP+ cells were found spread through the body of the animals, likely representing both resident and circulating macrophages as well as microglia cells in the brain (Figure 1A, Supplementary movie S1). Like macrophages and microglia in other species, morphologically the eGFP+ cells showed a variety of shapes, including spherical, amoeboid and dendritic (Figure 1B). Notably, macrophages in many species are notoriously auto fluorescent [37–39]. To ensure that the endogenous eGFP signal was not a result of autofluorescence, we performed immunostainings against GFP. We observed 100% colocalization between endogenous GFP and antibody-derived GFP signal, indicating that the endogenous fluorescence indeed corresponds to the expression of the transgene (Supplementary Figure 1 A-C, P). To determine whether macrophages are specifically labelled in different tissues in our mpeg1:GFP transgenic line, we tested the colocalization of eGFP+ cells with an established macrophage marker, colony-stimulating factor 1 receptor (CSF1R), in eye tissues. We observed eGFP+ cells and CSF1R colocalization in the cornea, vitreous chamber, and iris stroma of newt eyes (Figure 1C). We then performed immunostainings against two other well-established markers of macrophage populations, F4/80 and L-plastin. F4/80 is a glycoprotein expressed on the cell surface of several subtypes of mature macrophages such as microglia, Langerhans cells, and resident populations in the heart, kidney, and connective tissue [40]. This marker is not expressed on the cell surface of all macrophages and the levels of antigen expression differ depending on the environment in which the macrophage is found in mice [40]. In agreement with these studies [40], we observed that 41.4% of eGFP+ cells from several tissues in the newt body (tail, trunk, head) were F4/80+, indicating that this model allows identification of mature macrophages (Supplementary Figure 1 D-I, P). L-plastin is an actin-bundling protein also expressed by macrophages [41]. Notably, we also observed that a significant fraction of the eGFP+ cells (21.6%) co-express L-plastin, further indicating that this transgenic model enables labeling of macrophage populations (Supplementary Figure 1 J-O, P). Furthermore, we tested *in vivo* the phagocytic nature of the eGFP+ cells by examining the ability of eGFP+ cells to phagocytize glucan-encapsulated siRNA particles (GeRPs). GeRPs have been shown to be selectively incorporated by macrophages in several animal models and tissues [42–44]. We first injected GeRPs containing rhodamine in the cerebrospinal fluid through intraventricular injection. Right after the injection, we noticed the first particles being approached and encapsulated by eGFP+ cells in the central nervous system (CNS) (Figure 1D, Supplementary Movies S2, S3). The tissue analysis 20 hours post-injection showed that GeRPs had been phagocyted by microglia and border associated macrophages along the CNS (Figure 1E). When we injected GeRPs intraperitoneally (Figure 1D’), we found that, consistent with the previous experiment, GeRPs had been phagocytized locally by eGFP+ cells situated in the intraperitoneal cavity (Figure 1F).

**Figure 1.**
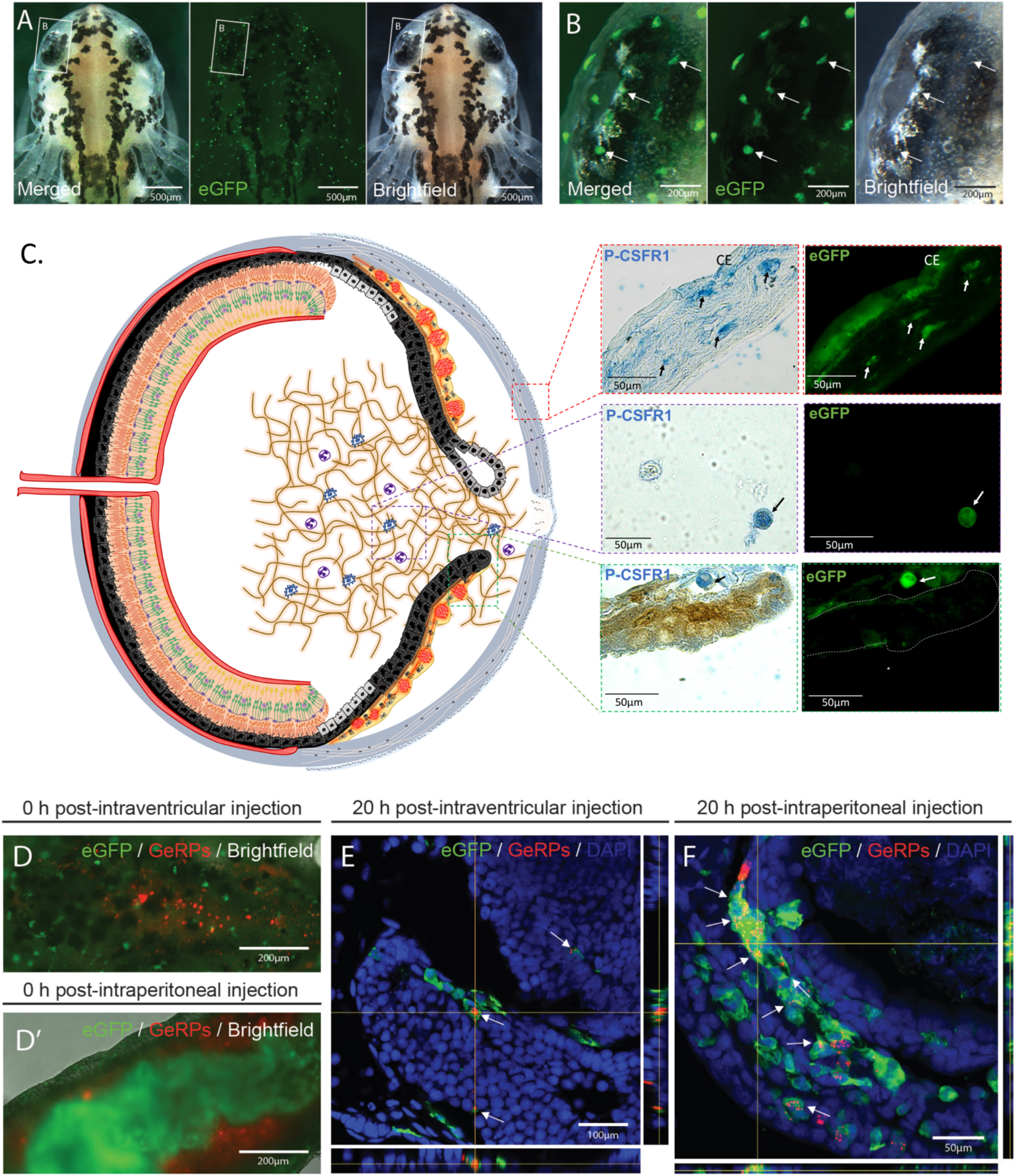
*mpeg1:GFP* transgenic newts enable the *in vivo* labeling of macrophages. **(A)** Representative images of an mpeg1:GFP+ initial larva (Developmental Stage 34) showing the widespread distribution of eGFP+ cells. Scale bar: 500 µm; n=3. **(B)** Image of the initial larval eye showing the region indicated in (A) at higher magnification. Arrows point to mpeg1:GFP+ cells showing spherical, amoeboid and dendritic morphology. Scale bar: 200µm **(C) (Right)** Double staining (paraffin embedded tissue) for eGFP and the macrophage marker P-CSFR1 in eye tissues such as cornea, iris stroma and vitreous chamber. Arrows highlight cells positive for both markers. Scale bars: 50µm. **(C) (Left)** Schematic drawing of the newt eye at 4 days post-lentectomy (dpl) (Stage 0-I). **(D)** Image of the dorsal view of the brain area of an mpeg1:GFP larva showing successful intraventricular injection of glucan-encapsulated siRNA particles (GeRPs). Scale bar: 200µm **(D’)** Image of the ventral view of the abdominal region of an mpeg1:GFP larva showing successful intraperitoneal injection of GeRPs. Scale bar: 200µm. **(E)** The vast majority of GeRPs were found phagocyted inside the eGFP+ cells (arrows) in the brain 20 hours post-intraventricular injection (O.C.T./Cryo embedded tissue). Scale bar: 100µm. **(F)** Similarly, the eGFP+ cells located in the intraperitoneal cavity were able to engulf most of the GeRPs (arrows) 20 hours after intraperitoneal injection (O.C.T./Cryo embedded tissue). Scale bar: 50µm.

Altogether, these results show that the new mpeg1:GFP *Pleurodeles waltl* transgenic line labels phagocytic macrophages *in vivo*.

### Macrophages accumulate transiently in the newt eye after lentectomy

To characterize the spatiotemporal recruitment of macrophages after lens removal, we analyzed eyes from mpeg:eGFP transgenic animals. Samples were collected prior to lentectomy, at 6 hours post-lentectomy (hpl), and 4-, 10-, 15-, and 30-days post-lentectomy (dpl). Few eGFP + cells were observed in the intact eyes, with most of them located at the corneal epithelium (Figure 2A i). At 6 hpl, eGFP+ cells were found in the cornea epithelium and inside the blood vessels of the iris (Figure 2A ii). At this time, the slit that was made in the cornea during the surgical removal of the lens was not yet closed. By 4 dpl, most macrophages were found around the wound area of the cornea and in the anterior eye chamber near the dorsal and ventral irises (Figure 2A iii). By 10 dpl, the corneal incisions were closed, and most macrophages were located near the regeneration-competent dorsal iris (Figure 2A iv). At 15 dpl, once the lens vesicle had formed, macrophages were located in the aqueous chamber (Figure 2A v). By 30 dpl, very few macrophages were detected in the anterior eye chamber (Figure 2A vi). Our data suggest that the macrophage mediated innate immune response to lentectomy in newts is resolved after regeneration, through which increased number of macrophages transiently populate the different anatomical structures of the eye.

**Figure 2.**
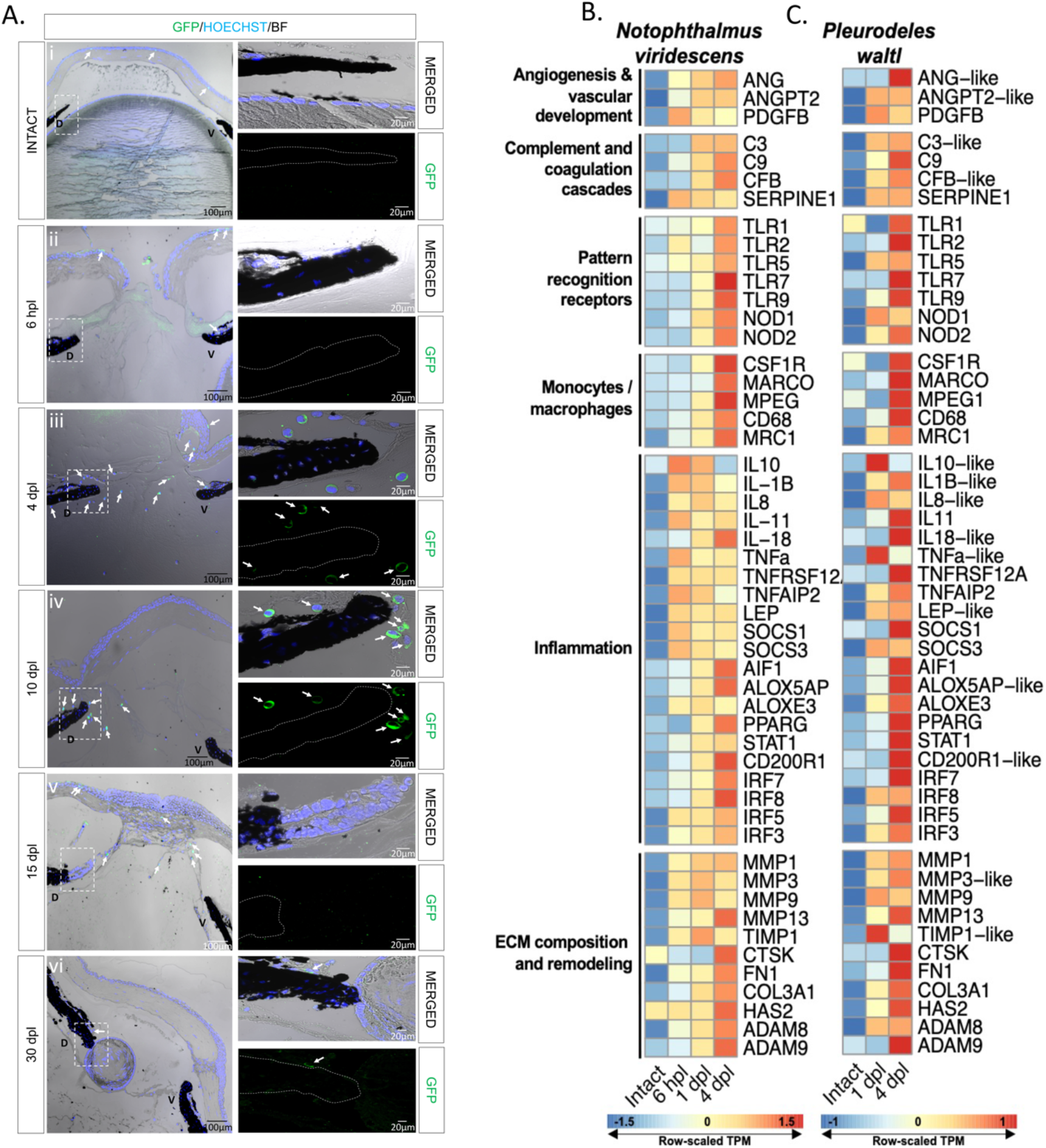
Visualization of macrophage recruitment and identification of highly enriched immune transcripts during lens regeneration. **(A)** Spatiotemporal view of macrophage recruitment in the newt eye during different stages of lens regeneration. eGFP immunohistochemistry (paraffin embedded tissue) was performed on eyes collected from mpeg1:GFP transgenic newts from intact eyes, 6 hpl (Stage 0) and 4 (Stage 0-I), 10 (Stage I-II), 15 (Stage III-IV)and 30 (Stage VIII) dpl. Macrophages were detected in the cornea of intact eyes and around the dorsal (D) and ventral (V) iris during the early stages of lens regeneration (arrows); n=6. Scale bars: 100µm (overviews, left) and 20µm (insets, right). **(B, C)** Heatmap displays the row-normalized expression levels of select immune-related transcripts in the dorsal iris of *Notophthalmus viridescens* (B) and of *Pleurodeles waltl* (C) following lens removal. The shown transcript identities were assigned by blastx annotation.

### Lens removal triggers a complex early response with a strong immune signature

Wound healing involves distinct phases that are well characterized at the molecular level in mammals. To characterize this highly complex response in newts, especially with regard to lens regeneration, we performed bulk RNA sequencing on two newt species (*Pleurodeles waltl* and *Notophthalmus viridescens*). We investigated dorsal irises from uninjured eyes, as well as irises following lentectomy at early time points corresponding to wound healing phases of lens regeneration. Under homeostatic conditions, the vertebrate eye is considered to be an immune-privileged organ due to several structural features and immunomodulator mechanisms that act together to limit inflammation [45–47]. As anticipated, prior to lens removal, we observed relatively low expression of immune-related transcripts in both newt species (Figure 2B&C). Lentectomy triggered a complex immune response, indicated by the upregulation of transcripts involved in the activation of signaling pathways pertaining to inflammation, ECM remodeling, pattern recognition, macrophages/monocytes, vascular development, complement activation and angiogenesis (Figure 2B&C). Consistent with the macrophage dynamics we observed from the mpeg1:eGFP line, RNA sequencing revealed a marked upregulation of macrophage related transcripts at 4 dpl (Figure 2A-C). A time course differential expression analysis in *Notophthalmus* identified time-dependent regulation of homologs associated with the KEGG pathways: TNF Signaling Pathway, Toll-Like Receptor Signaling Pathway, Inflammatory Mediator Regulation of TRP Channels, and ECM-Receptor Interaction (Supplementary Figure 2). We observed the significant up-regulation of several transcripts exhibiting homology to well-known pro-inflammatory cytokines such as interleukin-1 beta (IL-1β) and tumor necrosis factor alpha (TNF-a), as well as anti-inflammatory cytokines such as interleukin-10 (IL-10) between 6 and 24 hpl in both species (Figure 2B&C, Supplementary Figure 2 A&C). These observations are consistent with previous reports that show an early upregulation of anti and pro-inflammatory transcripts in regeneration competent species [12, 15, 16]. By 4dpl, the expression of these cytokines was reduced in *Notophthalmus viridescens*, and transcripts encoding proteins that can function as inflammatory mediators such as CD200R, AIF1, ALOX5AP, ALOXE3 and PPARG were upregulated in both species. Transcripts encoding protein products resembling ECM components, such as collagens (COL3A1), fibronectin (FN1), and hyaluronic acid (HAS2), were also upregulated following lens removal (Figure 2B&C, Supplementary Figure 2D). In addition, we observed the upregulation of transcripts that exhibited high homology to effectors of ECM remodeling, such as MMPs, ADAMs, and TIMP1 (Figure 2B&C). Our bulk RNA sequencing data paves the way to a better understanding of a genetic response to lentectomy that triggers lens regeneration in newts.

### Macrophage depletion inhibits regeneration by affecting cell cycle re-entry of iPECs

To test the role of macrophages during the early stages of lens regeneration, we injected control liposomes or clodronate liposomes into the eyes of the two newt species (Figure 3A). Liposomes are phagocytized by macrophages and, if carrying clodronate, ultimately induce apoptosis [48, 49]. Lens regeneration was evident by 30 dpl in control treated animals, but not in macrophage-depleted eyes, as indicated by the absence of a lens and ⍺A-Crystallin (a lens-specific marker) in both species (Figure 3B, C). In addition, histological assessment revealed several morphological and cellular abnormalities in the eye and near the dorsal and ventral aspects of the iris. Furthermore, an unusual cellular accumulation in the anterior and posterior eye chamber was observed, resembling the formation of scar-like tissue (Figure 3B, C). The inhibition of lens regeneration, morphological alterations, and cellular accumulation phenotypes were observed in 100% of the cases tested (n=40 per species). These data indicate that macrophages are essential to achieve lens regeneration in newts, as their depletion leads to the formation of scar-like tissue instead of the formation of a new lens.

**Figure 3.**
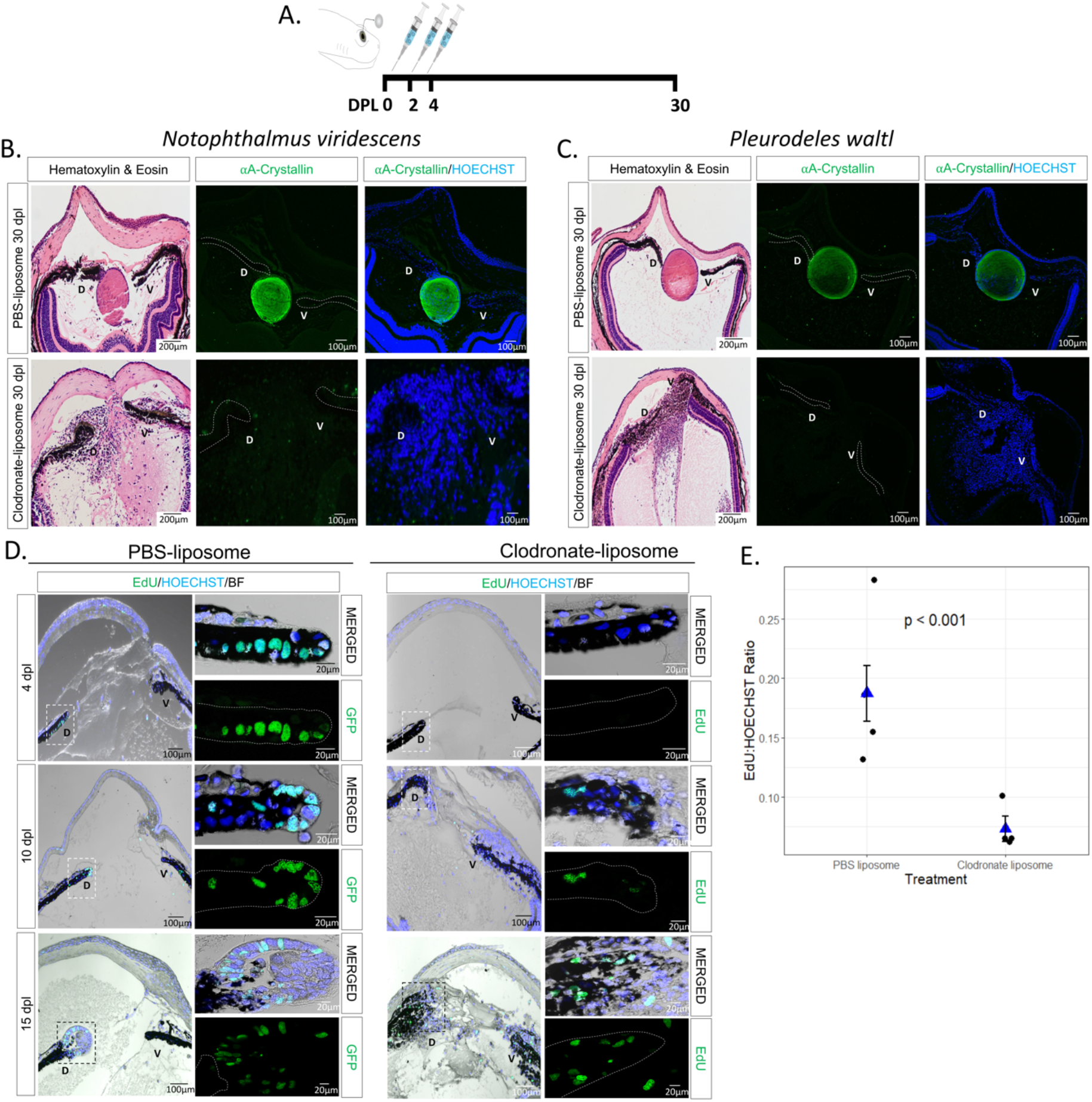
Early macrophage depletion inhibits lens regeneration and inhibits cell cycle re-entry of iPECs. **(A)** Schematic representation of experimental design. PBS or clodronate liposomes were administered intraocularly following lentectomy at 0, 2 and 4 dpl. **(B)** Lens regeneration in *Notophthalmus viridescens* was inhibited following early macrophage depletion indicated by the absence of ⍺A-Crystallin staining (paraffin embedded tissue). In addition, cellular accumulation was observed in the eye cavity at 30 dpl (Stage VIII); n=40. The dorsal (D) and ventral (V) iris epithelium were marked with dashed lines to aid in visualization. **(C)** *Pleurodeles waltl* displayed a similar phenotype following intraocular administration of clodronate liposomes; n=40. Scale bars: 200µm (Hematoxylin & Eosin, left) and 100µm (immunostainings, right) (paraffin embedded tissue). **(D)** EdU assay performed on eyes collected at 4 (Stage I), 10 (Stage II-III) and 15 (Stage IV-V) dpl following PBS and clodronate treatment (paraffin embedded tissue). The dorsal (D) and ventral (V) iris epithelium were marked with a dashed line to aid in visualization. Inset images of the dorsal iPECs highlight the effects of macrophage depletion on cell cycle re-entry; n=4 per time point. Scale bars: 100µm (overviews, left) and 20µm (insets, right). **(E)** Quantification of differences in the ratio of cells entering the S phase between clodronate and PBS treatment at 4 dpl; n=4. We estimated that the ratio of EdU to HOECHST cells was 0.187 (standard error 0.023) for the PBS liposome condition, while the ratio was 0.073 (standard error 0.011) for the clodronate liposome condition. Negative binomial regression, p<0.001.

Next, we sought to explore potential mechanisms by which macrophage depletion could inhibit lens regeneration. Previous work suggested that macrophage depletion affects the survival of progenitor cells during fin regeneration in zebrafish [50]. We tested if similar mechanisms take place during newt lens regeneration by TUNEL staining. We found that macrophage depletion using clodronate liposomes did not lead to apoptosis of iPECs in the early stages of lens regeneration (up to 10 dpl; Supplementary Figure S3A). As reported previously, apoptotic nuclei were observed at basal levels in transdifferentiating lens epithelial cells (LECs) in control-treated animals once a lens vesicle is formed [51].

One of the most prominent events after lentectomy is the cell cycle re-entry of terminally differentiated iPECs [52]. We tested whether cell cycle re-entry is affected upon macrophage depletion by analyzing EdU incorporation in the lentectomized irises in *Pleurodeles waltl*. We observed that macrophage depletion changed the dynamics of cell cycle re-entry of dorsal iPECs (Figure 3D). Based on a Negative Binomial Regression analysis, we found strong evidence of differences (p << 0.05) in the log ratio of mean EdU+ cells to HOECHST+ cells at 4 dpl (Figure 3E). Note that we provide a justification in the Methodology section for use of Negative Binomial regression instead of a t-test; the t-test based analysis also yields p<0.05. In control eyes, cells located inside the lens vesicle and lens epithelium were EdU + at 10 dpl and 15 dpl respectively, whereas in clodronate liposome treated eyes, a lens vesicle failed to form, and EdU+ cells were detected inside the iris and inside the vitreous and aqueous chambers. We noticed that the regenerating lens appears bigger and more developed at 15 dpl in control treated compared to the mpeg1:GFP animals at the same time (compare PBS-liposome group, 15dpl in Figure 3D with 15dpl in Figure 2A). This observation is consistent with our previous study showing that lens regeneration is delayed in older *Pleurodeles waltl* [53]. Our data support that, while macrophage depletion does not cause an apoptotic response, macrophages may play a role in the induction of cell cycle re-entry of iPECs.

### Macrophage inhibition prolongs inflammation, alters ECM remodeling, and causes a fibrotic-like response

To characterize the effects of macrophage depletion on the ensuing inflammatory response after lentectomy, we performed RT-qPCR to measure changes in the expression of the pro-inflammatory cytokine, IL-1β. Consistent with our bulk RNAseq experiments (Figure 2B-C), IL-1β expression was upregulated after lentectomy and returned to basal levels by 30dpl in PBS liposome-treated eyes (Figure 4A). However, in clodronate-treated eyes, IL-1β expression was significantly higher than controls at 10 dpl and remained higher than controls even at 30 dpl (Figure 4A).

**Figure 4.**
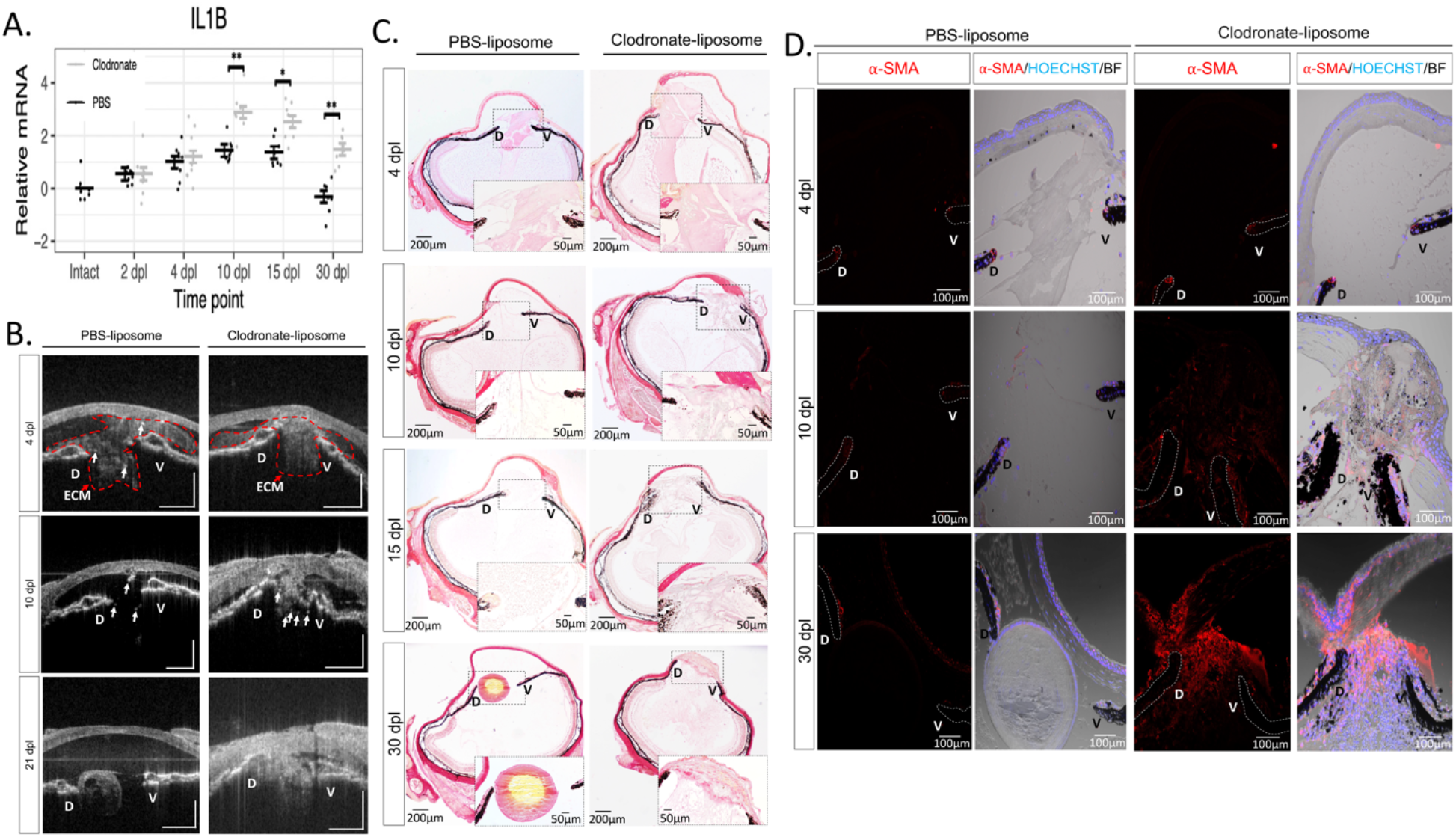
Early macrophage inhibition prolongs inflammation and disrupts ECM remodeling. **(A)** RT-qPCR analysis revealed an upregulation of IL-1β expression in clodronate-treated eyes at 10, 15, and 30 dpl; 8 eyes per treatment at each time point, *adjusted p < 0.005, **adjusted p < 0.0001. Error bars in plots represent standard error of mean estimate. Estimates were determined as described in methods to account for batch effects. Transcript abundance in the intact eye is shown for reference but was not included in the statistical analysis (see methods). **(B)** *In vivo* imaging of lens regeneration with SD-OCT shows the kinetics of ECM clearing in PBS liposome treated eyes. Following macrophage depletion, ECM remodeling was altered, and regeneration was inhibited; n=10. Arrows point to cloudy opacity in the SD-OCT image that is interpreted as ECM. **(C)** Deposition of collagen fibers were visualized with picrosirius red staining (Collagen stain red/pink in brightfield images) (paraffin embedded tissue). Collagen fibers were detected in the vitreous and aqueous chambers of PBS treated eyes at 4 dpl (Stage I) but cleared out by 10 dpl (Stage II-III); n=6. A progressive increase in collagen staining intensity was observed in clodronate treated eyes. Inset images show a higher magnification of the aqueous chamber. **(D)** The absence of macrophages triggers a fibrotic-like response. Myofibroblast presence was noted in the vitreous and aqueous chambers of macrophage depleted eyes at 10 (Stage II-III) and 30 dpl (Stage VIII), as indicated by ⍺-SMA staining; n=6. Scale bars: 100µm (paraffin embedded tissue).

We then used spectral domain optical coherence tomography (SD-OCT) to image the morphological changes that accompany macrophage depletion. The non-invasive nature of SD-OCT allows us to monitor the lens regeneration process from the same newt in real time [54]. Using SD-OCT, we have recently shown that ECM remodeling is highly correlated with lens regeneration [53]. At 4 dpl, cloudy opacity was observed in the anterior eye chamber between the dorsal and ventral iris of both PBS and clodronate treated eyes, indicating an accumulation of ECM (Figure 4B, Supplementary Figure S4). By 10 dpl, the opacity was cleared from the anterior chamber of control eyes and a newly formed lens vesicle was observed at the dorsal iris by 21 dpl (Figure 4B, Supplementary Figure S4, left panels). In contrast, ECM failed to clear and progressively worsened from macrophage-depleted eyes (Figure 4B, Supplementary Figure S4, right panels). Several other morphological abnormalities were detected in the newt eye by 21 dpl. The aqueous humor that fills the area between the cornea and iris failed to reform in clodronate-treated eyes, and bright spots were observed in the vitreous and around the dorsal and ventral irises (Figure 4B). We interpreted the bright spots as the cellular accumulation shown in Figure 3B-C. To confirm our interpretation of the SD-OCT findings, we used histology and picrosirius red staining to detect collagen fibers, a major component of the ECM [53]. Collagen accumulation appeared to be larger and denser in the anterior chamber of clodronate treated eyes compared to control eyes at 4 dpl (Figure 4C). In agreement with our SD-OCT interpretations, we observed that collagen fibers had begun to clear out at 10 dpl in control eyes and, by 30 dpl, very little staining was detected in the eye chambers. In contrast, collagen staining remained at 30 dpl in clodronate-treated eyes (Figure 4 C). Furthermore, we found that at least some of these cells are myofibroblasts, a cell type involved in ECM remodeling, immune modulation and angiogenesis [55] (Figure 4 D). To evaluate the kinetics of myofibroblast activation and accumulation in the anterior eye chamber, we performed immunostaining with alpha smooth muscle actin (⍺-SMA) at 4, 10, and 30 dpl in *Pleurodeles waltl*. While at 4 dpl we detected very few ⍺-SMA + cells in both experimental conditions, by 10 dpl, the accumulation of ⍺-SMA + cells was evident only in the anterior chamber of macrophage-depleted eyes, and this phenomenon was exacerbated by 30 dpl. In contrast, newts treated with PBS liposomes displayed substantially lower ⍺-SMA reactivity at 30 dpl relative to clodronate treatment (Figure 4D). Collectively, SD-OCT and histology demonstrated significant abnormalities following macrophage depletion in the newt eye including cellular accumulation and lack of ECM clearance, which resulted in the formation of scar-like tissue at the expense of lens regeneration.

### Exogenous FGF can rescue lens regeneration processes in macrophage-depleted newts

Previous studies have shown that FGF signaling pathway plays an important role during lens regeneration [56–60]. Since macrophages have been shown to directly secrete FGF in other contexts [61, 62], we hypothesized that macrophage depletion could affect the percentage of iPECs re-entering the cycle through a decrease in the FGF levels in the newt eye. To test this hypothesis, we supplied exogenous FGF2 into the newt eye right after lentectomy, followed by clodronate treatment (Figure 5A). We observed that FGF treatment caused a significant increase (p<0.001) in the overall number of EdU+ iPECs at 4 dpl (Figure 5B). Astonishingly, by 30 dpl, a lens vesicle was observed in one third of the cases, stemming from the dorsal iris of FGF2-supplemented eyes following clodronate treatment (4/ 12 eyes) (Figure 5C). In addition to rescuing regeneration, clodronate and FGF2 co-administration caused a reduction in the generalized cellular accumulation, as indicated by the lack of nuclear staining compared to clodronate and PBS co-administration (Figure 5C). Our data show that exogenous administration of FGF2 in macrophage-depleted newts can rescue lens regeneration as well as decrease the accumulation of cellular components of the scar-like tissue.

**Figure 5.**
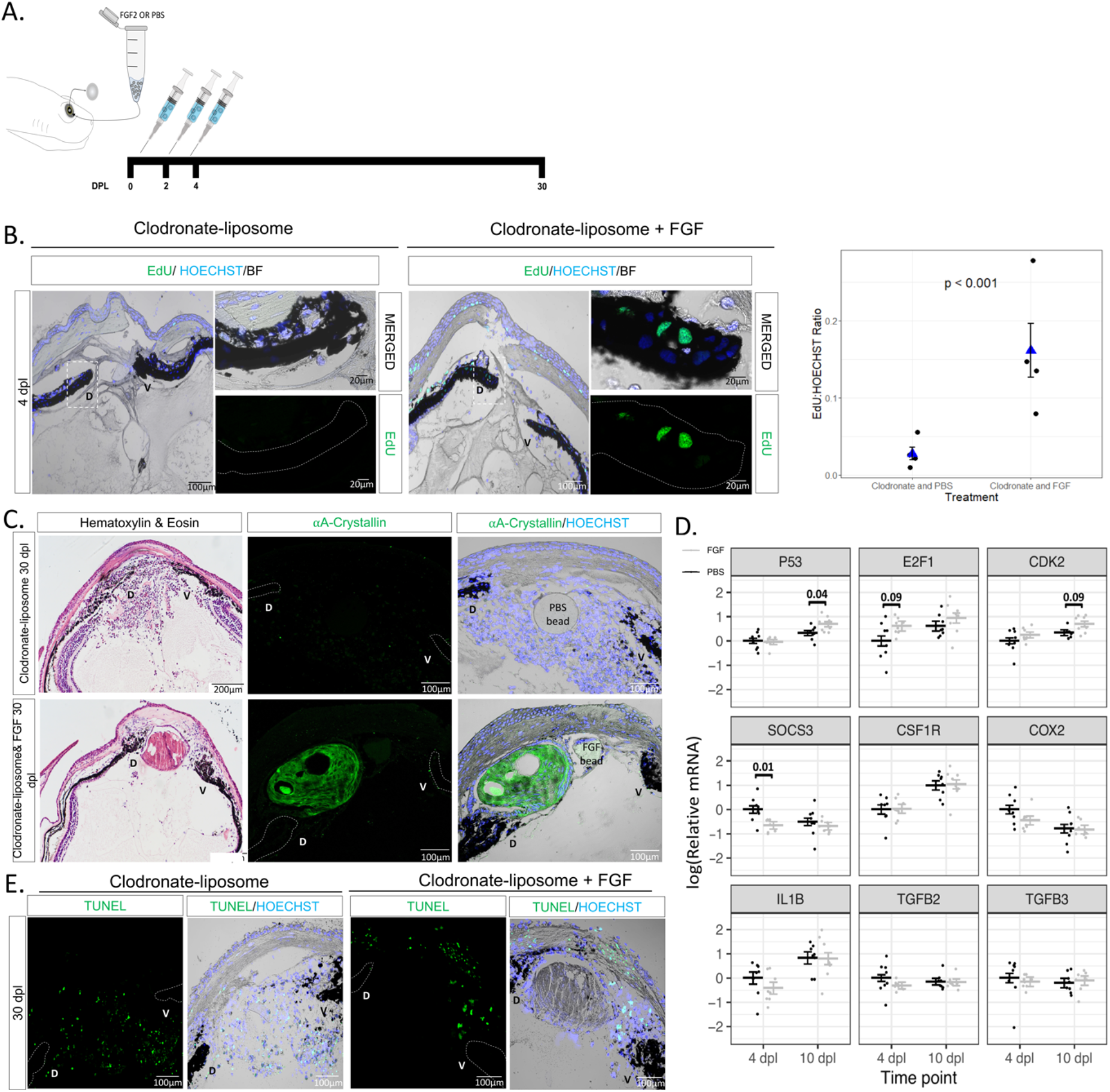
Exogenous FGF2 rescues cell cycle re-entry and lens regeneration following early macrophage depletion. **(A)** Schematic representation of experimental design. Clodronate liposomes were administered intraocularly following lentectomy at 0, 2 and 4 dpl, and a heparin bead incubated with PBS or FGF2 was added into the newt eye at 0 dpl. **(B)** EdU staining (paraffin embedded tissue) and quantification revealed a significant increase (p<0.001) in iPECs at the S-phase of the cell cycle in clodronate and FGF treated eyes; n=4. We estimated that the ratio of EdU to HOECHST was 0.161 (standard error 0.035) for the clodronate and FGF condition, while 0.028 (standard error 0.008) for the clodronate and PBS condition. Scale bars: 100µm (overviews, left) and 20µm (insets, right). Negative binomial regression, p<0.001. **(C)** Histology and immunostaining (paraffin embedded tissue) for lens specific marker ⍺A-Crystallin revealed that exogenous supplementation of FGF induced lens regeneration and resolve cellular accumulation at 20 dpl in 4/12 eyes. On the contrary, none of the 12 clodronate treated eyes that were supplemented with PBS beads had a crystallin lens; n=12. Scale bars: 200µm (Hematoxylin & Eosin, left) and 100µm (immunostainings, right). **(D)** RT-qPCR analysis for genes involved in cell cycle and inflammation at 4 and 10 dpl; 8 eyes per treatment at each time point. Statistical analysis using two-way ANOVA was performed and the adjusted p values displayed for p < 0.1. Error bars in RT-qPCRs plots represent standard error of mean estimate. Estimates were determined as described in methods to account for batch effects. **(E)** Detection of apoptosis in clodronate treated eyes with and without FGF supplementation. TUNEL+ nuclei were observed in the cornea and near the ventral (V) iris; n=12. Scale bars: 100µm (paraffin embedded tissue).

We next evaluated the gene expression levels of several inflammatory and cell cycle-related genes via RT-qPCR analysis at 4 and 10 dpl. The expression levels of cell cycle associated gene P53 was found to be significantly upregulated in clodronate/FGF2 treated eyes compared to clodronate/PBS treated eyes at 10 dpl. In addition, we observed moderate evidence (adjusted p < 0.1) that E2F1 and CDK2 were upregulated at 4 dpl and 10 dpl respectively in the FGF2-treated eyes. Furthermore, the expression of the suppressor of cytokine signaling 3 (SOCS3) gene was significantly downregulated at 4 dpl in clodronate/FGF2-treated eyes. In contrast, we did not find evidence for changes in the expression patterns of macrophage specific receptor CSF1R, inflammatory agents COX2, IL-1β, TGFβ2, and TGFβ3, (Figure 5D). We then tested if supplementing FGF in macrophage depleted eyes affected the apoptotic levels at 30 dpl. We found that apoptotic cells were still evident in the aqueous chamber near the regenerating lens and inside the cornea (Figure 5E). Our results suggest that FGF plays an important role during iPECs cell cycle re-entry and supplementation of exogenous FGF is sufficient to start the regeneration process even in the absence of macrophages.

### Late administration of clodronate enhances apoptosis and pro-inflammatory signals

To examine if newt macrophages play a role during the later stages of lens regeneration after the critical window of lens vesicle formation is passed, we treated eyes with clodronate or PBS liposomes at 10, 12, and 14 dpl (Figure 6A). Even though a lens was detected by 30 dpl, and LECs were positive for EdU and phospho-histone H3 (PHH3), the lens appeared smaller in all clodronate-treated eyes (Figure 6B). In addition to abnormal lens morphology (smaller size, presence of cells without fiber characteristics in the lens cortex and a multilayer lens epithelium), severe cellular accumulation was observed between the lens and the ventral iris (Figure 6B, C). Collagen fiber staining revealed that ECM accumulation was evident in the vitreous and aqueous chambers following clodronate treatments (Figure 6C). Furthermore, TUNEL+ nuclei were observed inside the lens fibers and near the ventral iris in clodronate treated eyes, indicating that macrophages may play a role in preventing cell death by apoptosis in the late phases of the regenerating lens (Figure 6D). Similar observations were noted when clodronate administration was initiated at 20 dpl (Supplementary figure S5). To complement, we tested for the expression of selected immune and inflammation related targets via RT-qPCR and found that late clodronate administration caused an increase in expression of the pro-inflammatory cytokine IL-1β and the macrophage-specific receptor CSF1R at 30 dpl (Figure 6E). Expression patterns of FGF2, TGFB2, TGFB3, MMP3, and MMP9 were not significantly changed (Figure 6E). Altogether, these data suggest that macrophages are also necessary for the proper regeneration of the lens in the later phases of the process, potentially controlling survival of lens cells and resolving pro-inflammatory signals.

**Figure 6.**
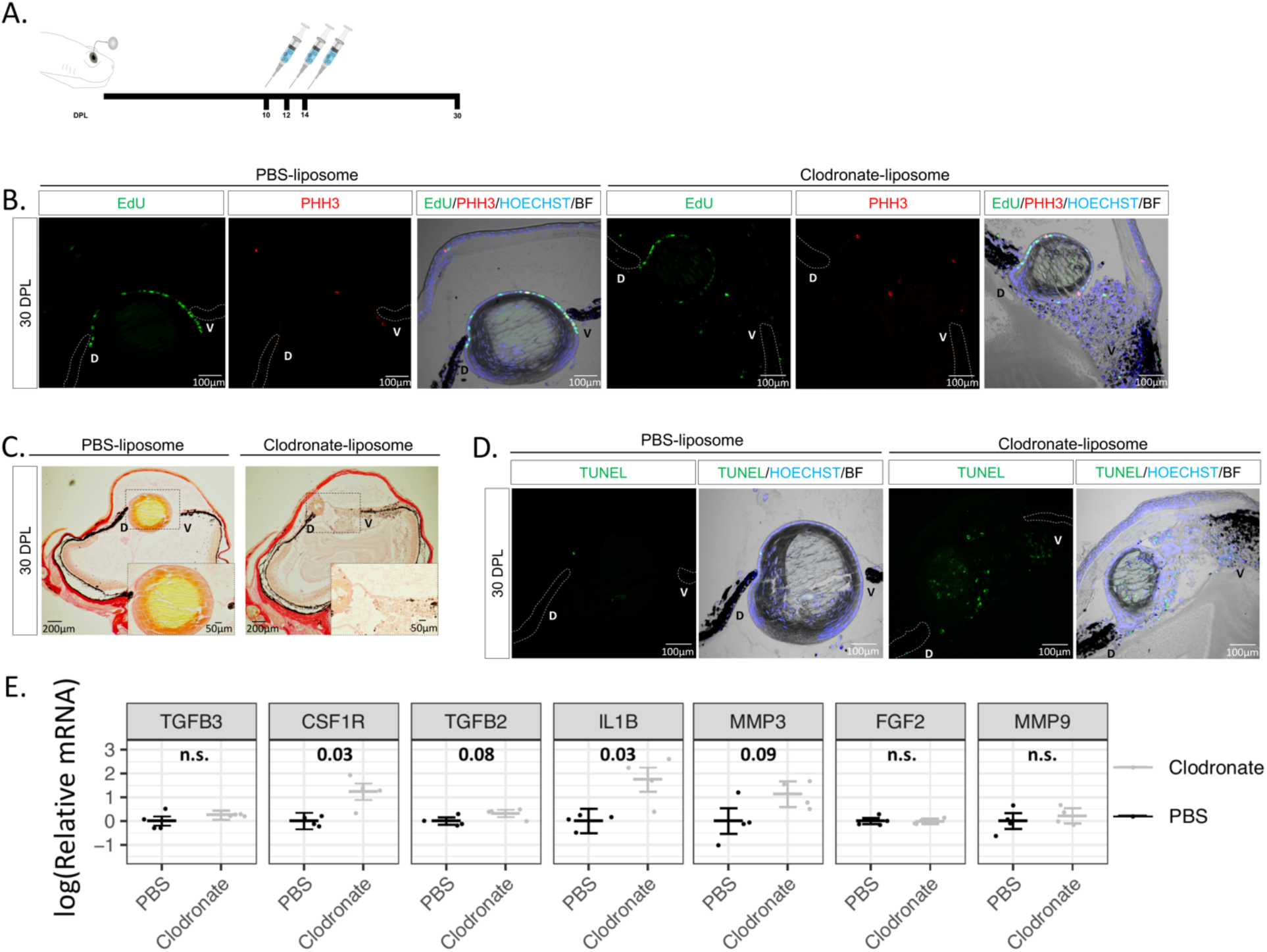
Late macrophage depletion after lens vesicle formation enhances apoptosis and pro-inflammatory signals. **(A)** Schematic representation of experimental design. Clodronate or PBS liposomes were injected intraocularly in the aqueous chamber at 10, 12, and 14 dpl after lens vesicle formation. **(B)** LEC in both treatments were positive for EdU and mitosis marker PHH3; n=6. Scale bars: 100µm (paraffin embedded tissue). **(C)** Collagen staining was more abundant in the vitreous chamber of the clodronate treated eyes; n=6. Inset images show higher magnification of the aqueous and vitreous chambers (paraffin embedded tissue). **(D)** Apoptotic cells were detected in the aqueous chamber and inside the regenerating lens; n=6. Scale bars: 100µm (paraffin embedded tissue). **(E)** RT-qPCR analysis revealed an upregulation of the anti-inflammatory gene IL1-β and macrophage receptor CSF1R following late clodronate administration; 4 eyes per treatment. Statistical analysis using Welch’s two-sample t-test was performed and adjusted p values displayed for p < 0.1. Error bars in plots represent standard error of mean estimate. n.s. = not significant

### A new injury helps to resolve the fibrotic phenotype and can re-activate lens regeneration

The results observed above led us to believe that macrophages in newts were playing an anti-fibrotic role during the injury response. This made us question whether macrophages, if given sufficient time to recover in number following clodronate depletion and lentectomy, could resolve the fibrotic injury that was triggered by their absence. To test this, we treated eyes with clodronate or PBS liposomes at 0, 2, and 4 dpl and monitored the animals using SD-OCT for 100 days, as well as H&E, EdU staining, and ⍺A-Crystallin immunohistochemistry, and we found no evidence of proliferation or lens formation at 100 dpl in all cases that were treated with clodronate (n=10) (Figure 7A). Startlingly, we also observed a microphthalmic phenotype in 5/10 eyes that were treated with clodronate (Figure 7B).

**Figure 7.**
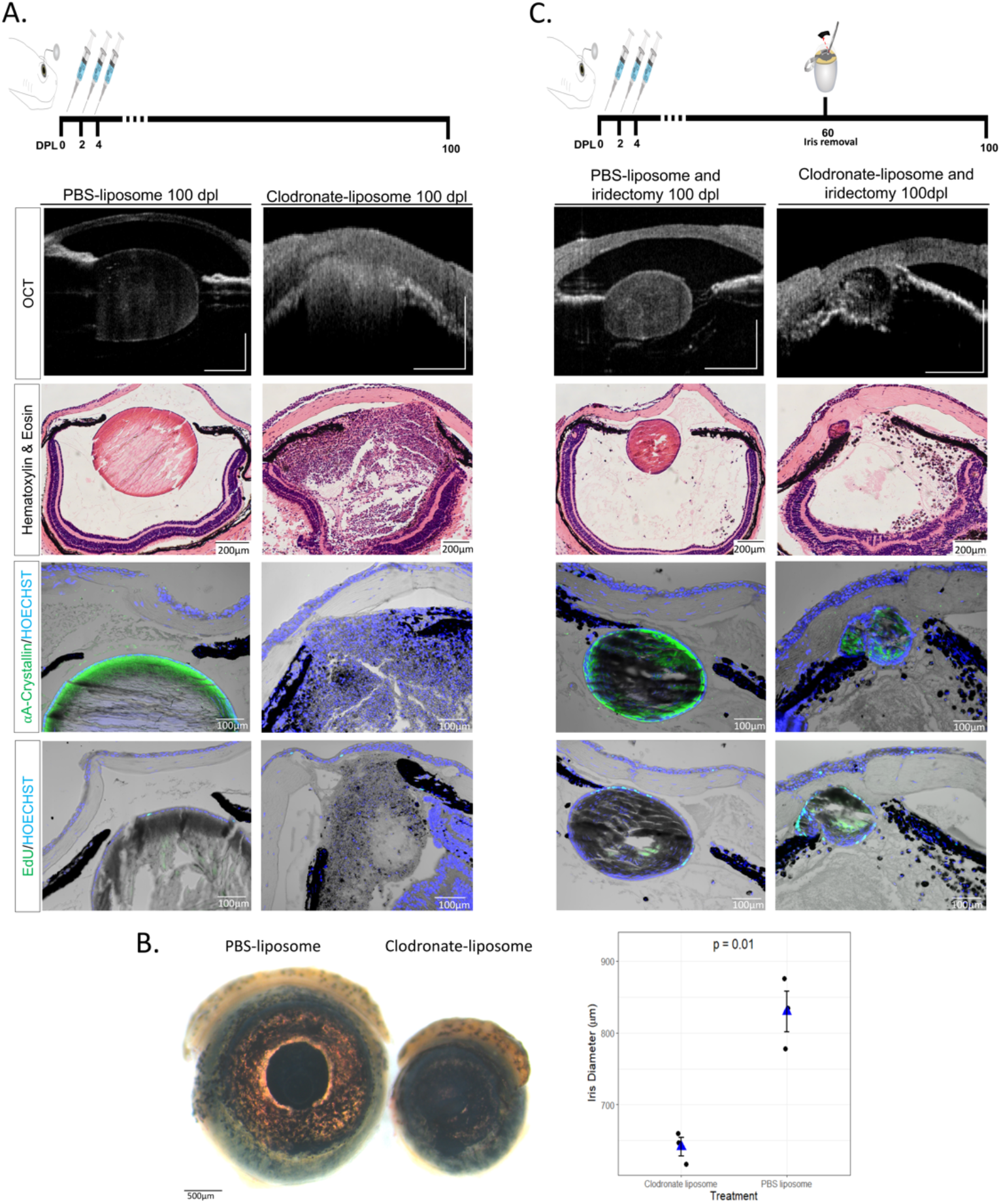
A secondary injury can induce the reabsorption of the scar-like tissue and re-initiate lens regeneration. **(A)** Schematic representation of the experimental design. Clodronate or PBS liposomes were administered at 0,2 and 4 dpl. At 100 dpl (Stage IX), 10 /10 clodronate liposome treated eyes showed severe cellular accumulation, and ECM (cloudy opacity) in the aqueous and vitreous chambers. Furthermore, no lens was observed in any of the clodronate liposome-treated eyes as indicated by the absence of lens specific marker ⍺A-Crystallin, and no EdU+ cells were observed in the lens (paraffin embedded tissue). On the contrary, 10 /10 of the PBS liposome-treated eyes regenerated a crystallin lens with EdU LECs without signs of ECM or cellular accumulation; n=10 per treatment. Scale bars from top to bottom: 100µm (OCT), 200µm (Hematoxylin & Eosin) and 100µm (immunostainings,). **(B)** At 100 dpl,microphthalmia was observed in clodronate treated eyes. Scale bar 500µm. Statistical analysis using Welch’s two-sample t-test was performed and adjusted p values displayed for p < 0.1. **(C)** Schematic representation of experimental design. Iridectomy was performed at 60 dpl and eyes were collected for histological analysis at 100 dpl. Secondary injury in the iris restarted the lens regeneration process, resolved cellular and ECM accumulation in 3 /8 clodronate liposome-treated and iridectomized eyes, as indicated by SD-OCT, histology, ⍺A-Crystallin and EdU staining at 100 dpl (40 days post-secondary injury); n=8 (paraffin embedded tissue). Regeneration was restarted in all PBS liposome-treated eyes; n=8. Scale bars from top to bottom: 100µm (OCT), 200µm (Hematoxylin & Eosin) and 100µm (immunostainings).

Since we observed a sustained inflammatory response with macrophage depletion, we hypothesized that the macrophages returning to the eye chamber were met with such an inflammatory microenvironment, that the macrophage phenotype was immediately polarized to a damaging, and hence continued, pro-inflammatory response upon entry into the eye. We wanted to test if a secondary injury was sufficient to reprogram the macrophages in order to resolve the chronic inflammation and fibrosis. To do that, we surgically removed a piece of the dorsal iris (iridectomy) at 60 dpl in PBS and clodronate-treated eyes without removing the fibrotic mass that was present in the anterior eye chamber (Figure 7C). It has previously been shown that following iridectomy, the iPECs proliferate to replace the missing tissue, and a lens is formed by the transdifferentiation of the newly formed iPECs [63]. Similar to these observations, we detected a newly formed lens following iridectomy in PBS-treated eyes (Figure 7C). Interestingly, we found that in 3/8 cases where iridectomy was performed in clodronate-treated eyes, regeneration was initiated, indicated by the presence of an ⍺A-Crystallin+ lens vesicle at the dorsal iris (Figure 7C). Importantly, the cellular accumulation and fibrotic phenotype was resolved from the anterior chamber in the cases that lens regeneration was induced (Figure 7C).

## Discussion

The requirement of macrophages for successful regeneration has been demonstrated in a wide range of vertebrates including axolotl, frogs, zebrafish, lizards, and African spiny mouse, but not in newts, the champions of regeneration, and never in the lens [11, 12, 15–20, 64, 65]. Here we used a newly generated transgenic newt line, mpeg1:GFP to show the dynamic recruitment of macrophages following lentectomy. Using bulk RNAseq data from two newt species, we uncovered upregulated expression of genes linked to macrophages/monocytes, inflammation, and other processes. We discovered that macrophage depletion during the early stages after lens removal inhibited regeneration in both *Notophthalmus viridescens* and *Pleurodeles waltl.* We went on to show that early macrophage depletion significantly reduced iPEC proliferation, but did not affect apoptosis. When we supplemented with exogenous FGF2, iPEC proliferation, the inflammatory profile, and regeneration were rescued. Furthermore, we showed that macrophage ablation failed to resolve the normal inflammation and ECM accumulation observed with injury, resulting in a deleterious eye environment which led to unresolved cellular accumulation, fibrosis, and eventually leading to microphthalmia. While a pro-inflammatory signature was detected when macrophages were depleted at later stages of regeneration, this experimental paradigm showed no changes in proliferation but presence of apoptosis. Importantly, we showed that re-initiation of an injury in the eye, broke the progressive cycle of inflammation, resolved the chronic fibrosis, and re-started the regenerative process. Collectively our data indicate that macrophages stand at a critical turning point during newt lens regeneration and determine the balance between scarring and regenerative phenotypes (Figure 8).

**Figure 8.**
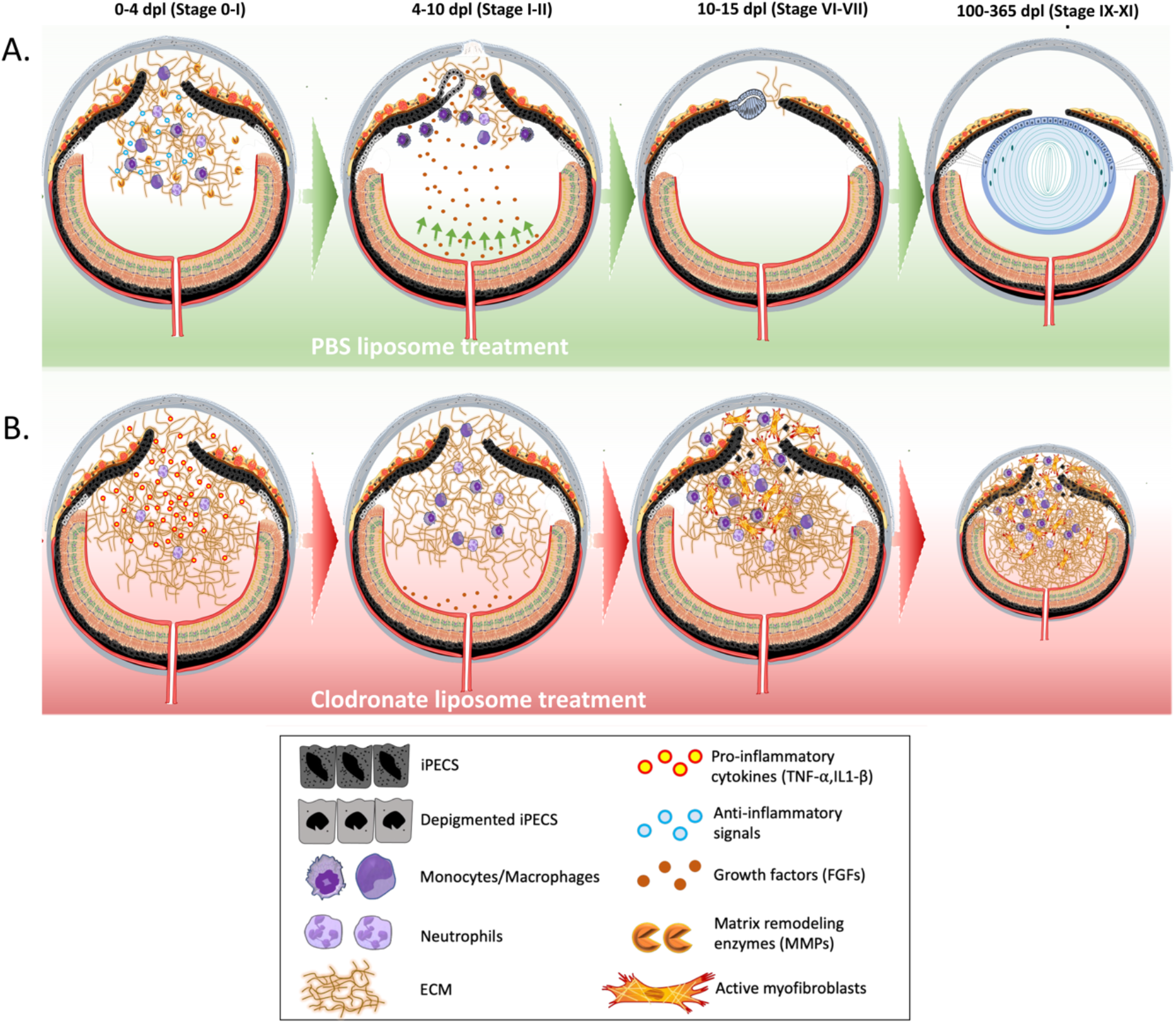
Simplified overview of the proposed model. **(A)** Following lentectomy, pro-inflammatory cytokines are secreted into the eye cavity and the anterior chamber of the newt eye fills with ECM. Macrophages and other immune cells accumulate in the eye by 4 dpl and secrete anti-inflammatory signals and matrix remodeling molecules to resolve inflammation and degrade ECM. Once ECM is cleared out, growth factors that are secreted from the neural retina reach the iPECs which then enter the cell cycle and start to dedifferentiate. During this phase, macrophages phagocytize the melanosomes that are discharged by iPECs [28, 31]. Dorsal iPECs become completely depigmented and give rise to a new lens vesicle. As LECs proliferate and differentiate into lens fibers, the regenerating lens becomes larger. **(B)** Upon clodronate treatment, the lack of anti-inflammatory cytokines and matrix remodeling molecules caused by the absence of macrophages at 4dpl, results in an increase and prolonged inflammatory state and the exacerbation of matrix accumulation. As a result, growth factors secreted from the retina can’t traffic through the damaged eye chamber and iPECs fail to re-enter the cell cycle. Alternatively, the macrophages are directly modulating the cell cycle re-entry of the iPECs. The pathogenic environment triggers the recruitment, differentiation, and activation of myofibroblast into the newt eye, contributing to the scar formation. Macrophages returning to the eye following clodronate treatment are unable to resolve the advanced inflammation and fibrotic environment. Absence of lens, severe fibrosis and chronic inflammation results in microphthalmia.

Despite being renowned for their regeneration capabilities, the utilization of newt species such as *Notophthalmus viridescens* and *Cynops pyrrhogaster* as model organisms has proven a difficult task [66]. Their gigantic genomes and complex life cycles have hampered the creation of molecular tools and the development of genetically modified animals. However, the application of recent technologies to *Pleurodeles waltl*, a newt species with similar regenerative capabilities [53, 66] which is easy to breed in the laboratory with a shorter reproductive cycle, and a recently sequenced genome, removed these roadblocks and brought newts into the molecular era [66–74]. Here, we utilized *Pleurodeles waltl* to develop the first macrophage reporter line in newts, using the macrophage-expressed gene 1 (mpeg1) promoter to drive the expression of *eGFP* [35, 36]. We validated the mpeg1:eGFP transgenic line by testing the colocalization of eGFP+ cells with known macrophage markers (F4/80, L-plastin, CSF1R) and demonstrating their phagocytic activity through local incorporation of GeRPs in several different tissues. We used this line to evaluate the spatiotemporal kinetic accumulation of macrophages in the newt eye during lens regeneration. Consistent with previous publications that used electron microscopy to characterize macrophage involvement during lens regeneration, we found a clear presence of eGFP+ cells between 4-10 dpl [34]. Interestingly, most of eGFP+ cells were near the regeneration-competent dorsal iris. The significance of this finding has not been evaluated in this report. However, future studies focusing on how the dorsal/ventral microenvironment affects the recruitment and phenotype of macrophages could be instrumental towards our understanding on the role of macrophages during scar-free healing. Nevertheless, the introduction of the mpeg1:eGFP transgenic line is a powerful addition to the regeneration toolbox and opens the door for a wide range of applications that were not before possible in newts.

It is hypothesized that the transition from wound healing to regeneration or scar formation is controlled by the complex interactions between immune cells, damaged tissues, and surrounding microenvironment [9, 75]. It is likely that the interaction between macrophages and iPECs during the early steps of wound healing provide critical signals for the initiation of cell cycle re-entry, dedifferentiation and transdifferentiation, as well as the resolution or propagation of inflammatory signals. We sought out to investigate this hypothesis and delineate the signaling pathways that occur during the early stages of wound healing in the newt eye via RNA sequencing. Our results showed an early upregulation of several genes associated with the resolution of inflammation, which is reminiscent to expression profiles reported in other regeneration-competent tissues, in contrast to a prolonged expression of pro-inflammatory signals that are dominantly observed in non-regenerative animals [12, 75–78]. In addition, we detected the upregulation of several matrix remodeling genes, including several members of the MMP family. The importance of MMP enzymes to degrade ECM and promote regeneration has been demonstrated in several species and tissues [18, 79, 80]. Furthermore, our data indicates the up regulation of genes in the lipoxygenase family. Lipid mediators play a key role in determining the magnitude and duration of inflammation in several species, however the role of these enzymes in regenerative animals is not yet well understood [81]. Overall, our RNA sequencing data suggest that during the early stages of lens regeneration, immune cells function to promote mechanisms that favor inflammation resolution and matrix remodeling and create a regeneration permissive environment. Further studies are required to specifically characterize the polarization state of macrophages and define the kinetics of M1 vs M2 class switching during lens regeneration in newts.

Macrophages can directly or indirectly regulate several aspects of the wound healing and regeneration processes. In zebrafish, for example, it was shown that macrophage depletion affects the survival of progenitor cells during fin regeneration [50, 82]. We tested if this is true during lens regeneration in newts and found no evidence of apoptosis in the iPECs of the dorsal iris at the early stages of regeneration in the absence of macrophages. We did, however, observe changes in the rate of cell cycle re-entry and proliferation of iPECs. Since macrophages can support proliferation through secretion of growth factors [83, 84], we tested if the addition of growth factors will counteract the effect of macrophage depletion in the newt eye. FGFs represent good candidates due to their known role in lens regeneration [56–59]. Astonishingly, when we exogenously supplemented a source of FGF in the absence of macrophages, the cellular accumulation was resolved and iPEC proliferation and regeneration were recovered. In addition, we showed that expression of *SOCS3* was downregulated in eyes that were supplemented with FGF2. Inhibition of SOCS3 has been shown to promote liver and axon regeneration in mammals [85, 86]. How macrophages either directly or indirectly affect the expression of *FGFs* in the newt eye requires further exploration. One possible explanation is that macrophages directly secrete FGF into the iris to promote proliferation [62, 83]. It’s also possible that macrophage depletion indirectly affected FGF levels by preventing its trafficking from nearby tissue sources. It has been documented, that during lens regeneration, the neuroretina secretes growth factors that are necessary for the reprogramming of iPECs [87, 88]. We showed that lentectomized eyes treated with clodronate liposomes exhibited higher levels of collagen in the aqueous and vitreous cavities. Therefore, it is possible that excessive amounts of ECM accumulation in the eye cavity could negatively affect the trafficking of growth factors from the neuroretina into the iPECs [89, 90].

Several studies have demonstrated that initiation of inflammation is necessary for successful regeneration [50, 91–95]. However, the magnitude and duration of inflammation is also key for determining the wound healing outcome [96, 97]. It is hypothesized that resolution of inflammation occurs much faster in regeneration-competent animals compared to non-regenerative species such as mammals [98, 99]. In fact, pharmacological attenuation of inflammation promoted tissue repair in regeneration-incompetent animals, thus demonstrating the importance of resolving inflammation during the early stages of wound healing [12, 100]. Increasing evidence suggests that macrophages are responsible for placing the necessary restraints in the inflammatory reaction during the early stages of regeneration [15, 16, 50, 101]. Consistent with these studies, we show that macrophage depletion in the newt eye prolonged the increased expression of the pro-inflammatory cytokine IL1-β.

In mammals, prolonged expression of inflammatory genes often leads to fibrosis and ultimately scarring [102–104]. This phenomenon is exacerbated during aging and repeated damage [105, 106]. Salamanders, on the other hand, have mastered scar-free healing and are thought to be resilient to fibrosis. This ability was highlighted in a pioneering study, where the lens was removed 18 times from the same newts in a period of 19 years, and each time, the lens was perfectly regenerated without any signs of fibrosis or scarring [107, 108]. It is tempting to speculate that the resistance to fibrosis in these animals could be due to the fact that newt macrophages adopt an earlier and more robust anti-inflammatory phenotype relative to their mammalian counterparts. It is also possible that newt macrophages can directly secrete collagens, MMPs, or growth factors that would modulate myofibroblast activation and shape extracellular matrix remodeling [18, 62, 109–111]. We tested the kinetics of ECM remodeling during normal lens regeneration and after early macrophage depletion with clodronate. Using our SD-OCT imaging modality in combination with histology, we detected the unprecedented formation of scar-like tissue in the newt eye following lentectomy with macrophage depletion and that ECM failed to clear. In addition, collagen composition and myofibroblast activation was altered with macrophage depletion. Our findings suggest that under normal conditions, newt macrophages play a critical role in preventing myofibroblast accumulation and fine-tune matrix remodeling. Defining the composition of ECM during injury in newts and identifying the complex interaction between ECM and immune cells that lead to its breakdown will be beneficial towards our understanding of ECM involvement during regeneration.

Interestingly, depletion of macrophages during late stages of regeneration, after the lens vesicle has formed, did not affect proliferation but it increased apoptosis in the lens fibers and around the ventral iris. Clodronate treatment also arrested the regeneration process, induced expression of the pro-inflammatory cytokine IL1β, and resulted in unresolved cellular accumulation. Our findings suggest that macrophages also play important roles after the critical window of transdifferentiation: they contribute to promote and maintain homeostasis in the eye by reducing the pro-inflammatory environment, preventing scar formation and keeping apoptosis at bay.

Surprisingly, when we monitored clodronate-early-treated eyes for 100 dpl, we found that the scar-like tissue components (cellular inflammation, ECM accumulation, and fibrosis) progressively worsened and, in most cases, microphthalmia was observed. These observations are interesting because macrophage depletion with clodronate is transient, yet the return of macrophages appeared unable to resolve the ECM accumulation that occurred in the absence of macrophages at the key early timepoints. It is unclear whether the macrophages returning to the fibrotic lesion and altered microenvironment have lost the ability to regulate ECM dynamics or whether the ECM has progressed into a state that the returning macrophages are unable to modify. Astonishingly, introduction of a secondary injury without removing the scar tissue, by removing a small piece of the dorsal iris was sufficient to resolve the deleterious effects of macrophage depletion and re-start the regeneration process. These findings are noteworthy and suggest that signaling events are capable of reprogramming newt macrophages into a pro-resolving and anti-fibrotic phenotype, even in the presence of a fibrotic environment. Eliminating myofibroblasts has been a long-standing goal for developing therapies to treat fibrosis in humans. In newts, however, macrophages can achieve this when activated in response to a new injury. Even more impressively, they can do this after chronic inflammation and at a point which stable fibrotic disease has already been established [109]. Consistent with our observations, Godwin et al., have previously shown that re-amputation of fibrotic limbs after macrophage depletion in axolotls resulted in regeneration rescue [16]. However, in these experiments the entire fibrotic environment was removed by amputation, whereas in our study, we kept the vitreous and aqueous chambers (area filled with collagens, cellular accumulation and myofibroblast) and only removed the source of regeneration (dorsal iris epithelium) accentuating the significance of our findings. Our findings illustrate that newts can not only re-start a previously hampered regeneration process, but they are able to correct previously established fibrotic tissue when exposed to a new insult. Future studies comparing the phenotype of newt macrophages during normal lens regeneration in control animals with those recruited to the fibrotic environment after clodronate treatment and those after secondary injury has potential therapeutic interest, as this approach could give us clues to which polarizing factors are responsible for inducing anti-fibrotic and pro-regenerative outcomes in newts.

There are several limitations in the present study. First, bulk transcriptomic methods were used to characterize the molecular mechanisms that promote wound healing and inflammation resolution during the early stages of regeneration. The iris epithelium and stroma consist of a heterogeneous cell population (iPECs, keratinocytes, melanocytes, muscle, blood, and infiltrating immune cells such as macrophages). In axolotls and zebrafish, it was shown that not only immune cells, but also other cell types such as blastema progenitors and fin epithelial cells (in limb and fin respectively), can also produce cytokines that modulate the inflammatory response and influence the regeneration outcome [50, 112]. Future studies utilizing technologies with the ability to resolve cell types, such as cell sorting, single-cell RNA sequencing, and/or spatial transcriptomics will shed light on how each cell population contributes differently during lens regeneration. Furthermore, these technologies will allow us to identify the heterogeneity of the cellular accumulation that we observed following clodronate treatment. Another limitation comes from the use of liposomes to deplete macrophages. Since liposomes cannot penetrate the blood-brain barrier, they must be injected intraocularly, thus creating an additional injury to the eye. Pharmacological studies using small molecules to directly target macrophages and the generation of animals with genetically depleted macrophages will aid in further exploration of the function of macrophages during scar-free healing in newts.

Collectively, our results suggest that lens regeneration in newts can be added to the ever-growing list of cases that demonstrate the necessity of macrophages for successful regeneration. Identifying the common mechanisms by which macrophages contribute to scar-free healing across different regenerative species and tissues will be critical towards our goal to integrate these findings for clinical translation and the development of therapeutic strategies. This study offers a unique perspective and a first glimpse into the functions of macrophages to achieve successful regeneration in the newt eye: by promoting early cell cycle re-entry in iPECs, resolve pro-inflammatory signals, and prevent fibrotic scar formation. Furthermore, we showed that macrophage depletion during lens regeneration establishes a progressive fibrotic disease state in the newt eye that could lead to microphthalmia. While we showed the fibrotic disease is unable to resolve on its own, a secondary injury performed months later in the dorsal iris resulted in the reabsorption of the fibrotic scar within the eye and the initiation of lens regeneration. The remarkable reversal of fibrosis with re-injury occurred in the presence of returning macrophages, highlighting their potential role in clearing the previously established fibrotic disease. Taking into consideration that fibrosis in humans is often considered irreversible, these observations are significant and bear great translational potential. Our findings establish a new experimental context in which the mechanisms behind scar-free healing, regeneration and scar reabsorption can be studied further.

## Methodology

### Animal husbandry and ethical statement

Iberian Spanish newts, *Pleurodeles waltl*, and red spotted eastern newts, *Notophthalmus viridescens* were used in this study. Spanish newts were bred and grown in our colony at Miami University, whereas red spotted eastern newts were wild caught. Handling and surgical procedures were performed following guidelines by the Institutional Animal Care and Use Committee at Miami University. Adult *Pleurodeles waltl* wildtype newts born and raised in captivity for generations were used for transgenesis in the aquatic facilities of Karolinska Institutet, Stockholm. Handling, breeding, transgenesis and the other experimental procedures performed in Stockholm were done according to both Swedish and European regulations. All animals were raised according to previously established husbandry guidelines [113].

It’s important to note, that the lens growth rates and speed of regeneration can vary between different newt species and between different ages of the same species [28, 52, 53]. For this reason, we refer to lens regeneration stages (as defined in [53]) in addition to days post-lentectomy throughout the manuscript.

### Generation of Tol2-mpeg1:eGFP-polyA plasmid

A vector containing a 1.86kb fragment of the zebrafish *mpeg1* promoter driving eGFP as previously described [35], was a kind gift from Enrique Amaya. The mpeg1 fragment was amplified by PCR (*mpeg1 F* 5’-TTGGAGCACATCTGAC -3’; *mpeg1 R* 5’-TTTTGCTGTCTCCTGCAC-3’) and subcloned into pBSII-SK-mTol2 upstream of the coding region of EGFP. SV40 polyA was used as termination signal. The resulting plasmid was purified using cesium chloride preparation to avoid potential contaminants that may negatively impact on transgenesis efficiency or newt survival. The plasmid was subsequently purified using Qiagen Maxiprep according to manufacturer’s instructions and then resuspended in ultrapure distilled water.

### *Pleurodeles waltl* transgenesis: tgTol2(Dre.*mpeg1:eGFP*)^MHY/SIMON^

Approximately 5nl of a mix composed of 1µl of Tol2-mpeg1:eGFP-polyA plasmid (100ng/ml) and 3µl of Tol2 transposase (300 ng/ml) was injected into single-cell eggs to generate transgenic founders (F_0_) adapting previously described procedures to *Pleurodeles waltl* [114]. The founder generation of mpeg1:GFP newts were bred with albino newts (Tyr^-/-^) [65] to establish the transgenic line. Every clutch from the first generation (F_1_) onwards was screened with a fluorescence microscope to select for the positive offspring (Figure 1A). Only animals from generation F_1_ onwards were used in this study.

### Phagocytosis Assay: Glucan-encapsulated siRNA particles (GeRPs) injections

GeRPs were a kind gift of Myriam Aouadi [42–44].1mg of GeRPs containing Rhodamine was resuspended in 1 ml of PBS and sonicated just before injection. The sonication protocol was: 40%, 10 sec; 35%, 10 sec; 30%, 10 sec; 25%, 10 sec; 20%, 10 sec; 30%, 30 sec. GeRPs were injected using pulled borosilicate glass capillaries (Harvard Apparatus, GC100F-10) in transgenic Mpeg1:eGFP *Pleurodeles* [F_1_: tgTol2(Dre.mpeg1:eGFP)MHY/Simon] either intraperitoneally or intraventricularly according to previous established protocol [115]. Live imaging of larvae and time lapse movies were created with Zeiss Axiovert 200 M inverted microscope. For histological observations, larvae were fixed at 20 hours post-injection and processed for immunofluorescence according to Joven et al., 2018 [70].

### Lentectomy, iridectomy and EdU injections

Animals were anesthetized by whole body submersion in 0.1% ethyl 3-aminobenzoate-methane Sulfonic acid solution (MilliporeSigma) diluted in amphibian phosphate-buffered saline (APBS :1x PBS plus 25% dH2O). Once the animal was anesthetized, a scalpel was used to make a slit in the cornea and the entire lens was carefully removed with fine tweezers [116]. Clodronate or PBS liposomes (Encapsula Nano Science, # CLD-8901) were injected intraocularly into the vitreous cavity of the newt eye using a pre-pulled glass needle (20μm tip diameter) attached to a microinjector (MicroJect 1000A, BTX, Harvard Apparatus) set at 10.5psi. For iridectomy experiments, the apical region of the dorsal iris (and regenerating lens from PBS-liposomes treated eyes) was surgically removed at 60 dpl by re-opening the cornea with a scalpel and removing a piece of the iris using scissors and tweezers. After all surgical procedures, the animals were allowed to recover from anesthesia and return to appropriate housing containers where they were monitored carefully for the duration of the experiments. For cell proliferation studies, EdU (Invitrogen, #C10338) was injected intraperitoneally 24 hours prior to collection at 10 µg/g of body weight. All animals used for these experiments were post-metamorphotic, juveniles at 6-8 months old.

### FGF2 experiments

Heparin-coated polyacrylamide beads (Sigma, #H-5263) were washed in APBS and incubated either with 0.25μg/μl bFGF (R&D Systems, #133-FB) or APBS (vehicle control) overnight at 4°C. Heparin beads were carefully inserted with tweezers into the vitreous chamber of the eye, between the dorsal and ventral iris following lens removal. All animals used for these experiments were post-metamorphotic, juveniles at 6-8 months old.

### *Notophthalmus viridescens* RNAseq, transcriptome assembly, and differential expression analysis

Dorsal iris tissues were collected from adult *Notophthalmus viridescens* from intact animals (no lentectomy), as well as animals 6 hpl, 1 dpl, and 4 dpl. Three biological replicates were used per time point, with each biological replicate containing bilateral dorsal irises from 3 or 4 animals pooled together. The iris tissues were placed in 500μL of cold TRI Reagent (Zymo, R2050-1-50), vortexed, and stored at -80 °C until RNA extraction. RNA isolation was performed via extraction with 0.2 volumes chloroform followed by processing of the aqueous phase with the Direct-zol RNA Microprep Kit (Zymo, R2060) according to manufacturer’s instructions, including an in-column DNase I treatment. RNA integrity was assessed with the Agilent RNA 6000 Pico Kit (Agilent, 5067-1513) and quantification performed with the Qubit RNA HS Assay Kit (ThermoFisher, Q32852). Reverse transcription and library preparation were carried out with the Zymo-Seq RiboFree total RNA library Kit (Zymo, R3000) using 108 ng input RNA per reaction. RiboFree depletion was applied for 4 hours to deplete overrepresented transcripts, and final libraries were amplified with dual indexes (Zymo, D3096) for 13 PCR cycles. Sequencing was performed at the Novogene sequencing core on the Illumina NovaSeq 6000 to approx. 100 million paired-end reads per sample.

The first 10 base pairs of sequence read2 were hard trimmed to remove low-complexity bridge sequences introduced during library preparation. The reads were then quality and adapter trimmed with Trim Galore using parameters *-q 5 --length 36 --stringency 1 -e 0.1* [117, 118]. rRNA reads were depleted by aligning trimmed reads against the Silva rRNA database using Bowtie2 [119, 120]. Cleaned reads were prepared for transcriptome assembly using the Trinity assembler pre-processing script insilico_read_normalization.pl with parameters *--seqType fq --JM 1450G -- max_cov 30 --pairs_together --SS_lib_type RF -CPU 24 --PARALLEL_STATS --KMER_SIZE 25 -- max_CV 10000 --min_cov 2* [121]. Trinity assembly was performed on each condition individually with Trinity using *parameters --seqType fq --max_memory 1450G --SS_lib_type RF --CPU 24 -- min_contig_length 200 --monitoring --min_kmer_cov 2 --no_normalize_reads*. Individual assemblies were merged into a final transcriptome with the DRAP pipeline runMeta tool and parameters *--strand RF --mapper bwa --length 200 --type contig --coverage 0,1,10 --write* [122]. Assembled transcripts were annotated against the NCBI non-redundant database using an implementation of blastx and the functional annotation module in the OmicsBox software environment [123]. Final transcript abundance was estimated with the Salmon alignment tool using the parameters *-l ISR --numGibbsSamples 20 --seqBias --gcBias --reduceGCMemory -d* [124]. Differential transcript abundance testing was performed in the R environment using the SWISH Fishpond workflow [125]. Expression values displayed in manuscript are the log-transformed Transcript Per Million (TPM) values, averaged across inferential replicates. In the case of transcripts with multiple detected isoforms, the transcripts displayed in the heatmap represent the transcript assigned the lowest E-value determined by blastx. KEGG pathway enrichment analysis was performed using the Combined Pathway Module inside the OmicsBox environment, with significance values calculated using maSigPro time-course analysis [126, 127]. For combined pathway analysis, parameters were set as *Keep most specific pathways =* TRUE and *Blast expectation value = 0.*001, and KEGG orthologs were linked through the EggNog mapper [128]. A two tailed Fisher’s test was applied to pathways and FDR cut off was applied at 0.05.

### *Pleurodeles waltl* RNAseq and differential expression analysis

Dorsal iris tissue was dissected from 13-month-old newts using intact or lentectomized animals. Tissue from 4 animals were pooled for each biological replicate, collected in triplicate. Samples were collected into cold TRI Reagent (Zymo, R2050-1-50), vortexed, and stored at -80 °C until RNA extraction, as described above. RNAseq libraries were prepared using 72-130 ng of isolated RNA, using NEBNext® Ultra™ II Directional RNA Library Prep with Sample Purification Beads (NEB, E7765S) and NEBNext® Poly(A) mRNA Magnetic Isolation Module (E7490S). Indexing was performed using NEBNext® Multiplex Oligos for Illumina® (NEB, E7500/7710/7730). Samples were pooled and sequenced on a lane of NovaSeq 6000 at the Novogene Sequencing Core using paired-end, 150 base pair reads to a minimum depth of 29 million read pairs per sample.

Reads were quality trimmed using Trim Galore with parameters *--stringency 3 --paired --length 36* [117, 118]. Cleaned reads were aligned to the *Pleurodeles waltl* genome using the STAR aligner with two-pass alignment to insert splice junctions [67, 129]. Aligned reads were assembled into transcripts using Stringtie and spliced transcripts were extracted using gffread [130, 131]. Assembled transcripts were indexed using the Salmon quantification tool, using the parameter *- k 31* and providing genomic sequence as decoy [124]. Transcript expression was quantified using salmon with parameters *-l ISR --validateMappings -p 12 --numGibbsSamples 20 --seqBias --gcBias -d*. Samples with alignment rates > 80% were included in downstream analysis, resulting in the exclusion of one sample (intact replicate 3). Expression values displayed in manuscript are the log-transformed Transcript Per Million (TPM) values, averaged across inferential replicates. Assembled transcripts were annotated against the NCBI non-redundant database using an implementation of blastx and the functional annotation module in the OmicsBox software environment [123].

### RT-qPCR design and analysis

Whole eyes were enucleated at the indicated time points and placed in 500μL of TRIzol and stored at -20 °C. The tissues were then homogenized mechanically using pellet pestles and centrifuged to remove debris. Total RNA was isolated using Direct-zol RNA Microprep (Zymo Research, #R2061) following manufacturer’s instructions. RNA yield and quality were analyzed using Nanodrop ND-2000 Spectrophotometer (Thermo Scientific) and Agilent 2100 Bioanalyzer (Agilent Technologies), respectively. cDNA was synthesized using 200 ng of total RNA as a template with QuantiTect Reverse Transcription kit (Qiagene, #205313) according to manufacturer’s instructions. The synthesized cDNA was diluted at 1:10 ratio with pure water. 2 ul of the cDNA dilution were used for quantitative PCR (qPCR) reactions. The final qPCR reactions contained: 2 ul of diluted cDNA, 10 ul of TB Green® Advantage® qPCR Premix (Takara, #639676) and 50 nM of each primer, adjusted to 20 ul with water. qPCR reactions were set up in duplicate in the Rotor-Gene Q thermocycler 5 plex (Qiagene, Germantown, MD, USA) using annealing temperature set at 60 °C. Primers reported here were designed using primer blast (https://www.ncbi.nlm.nih.gov/tools/primer-blast/) and obtained from IDT Technologies. The gene coding sequences were obtained from iNewt [132]. Primer and target sequences are shown in Table S1. TAII70, was used as a housekeeping gene using primers published previously [133]. The comparative Ct method was used to determine relative gene expression levels compared to the housekeeping gene. The primers and qPCR reactions were validated following qPCR MIQE guidelines [134]. Four to eight biological samples were used per condition.

In order to improve data conformance to statistical modeling assumptions, a natural log transformation was applied to relative mRNA values. For time course analysis of intact eyes through 30 dpl eyes (Figures 4A), a two-way block ANOVA was conducted, with treatment and time as the factors along with their interaction. The blocking factor was included to account for batch effects, as animals were collected in two groups. The analysis was done using the aov function in the R stats package [135]. Treatment differences were estimated for each time point using the emmeans function in R, and since five comparisons were made, the p-values were corrected by controlling the false discovery rate [136–138]. For experiments in which transcript expression was assayed at two time points (Figure 5D), a two-way ANOVA with interaction (factors Treatment and Time) was performed using the lm function in R, part of the stats package. Analysis of treatment means and multiple comparisons correction was done as described above. For the ANOVAs, residuals were examined to check for severe assumption violations, including constant error variance and normality. For experiments with a single time point (Figure 6E), a Welch’s two-sample t-test was applied using the R function t.test, part of the stats package. All animals used for these experiments were post metamorphotic, juveniles at 6-8 months old.

### Optical Coherence Tomography

The anterior chamber of each eye was non-invasively monitored via spectral domain optical coherence tomography (SD-OCT) as previously described [54]. The animals were anesthetized prior to imaging, and 100 µL of water was applied to the cornea surface with a pipette to reduce reflection artifacts caused by dehydration during live imaging. The broadband light source was filtered to produce a low-coherence spectrum center around 850nm with frequency range near 200 MHz, which can produce an axial resolution of 2 µm and lateral resolution of 8 µm. The scanning area of the eye was 1.2 by 1.2 mm, where 500 B-Scans were collected across this range. Each B-Scan consisted of 2000 A-Scan, where each A-Scan had 2048 pixels. The final C-Scans were rescaled and reconstructed as 500 x 500x 500 voxels. All animals used for these experiments were post metamorphotic, juveniles at 6-8 months old.

### Tissue embedding and sectioning

For cryo-embedding, collected tissues were washed 2 to 3 times in PBS and fixed overnight in 4% PFA (1 g paraformaldehyde in 25 mL PBS) at 4 °C. For cryoprotection, animals were transferred to new tubes with 30% sucrose (in 0.1M PBS) overnight at 4 °C. Tissues were placed in embedding molds, correctly positioned and tissue tek (yellow Shandon Cryochrome, Thermo Fisher Scientific, Waltham, USA) was gently added. Embedded molds were placed at -80 °C for 15 minutes and transferred to -20 °C and stored until sectioned. For sectioning, a cryostat was used with an object temperature of -21 °C and knife temperature of -19 °C. For experiments involving whole animals, 12 µm sections were made. Sections were collected on Superfrost Plus slides and stored at -20 °C. Tissue processing for the phagocytic experiments depicted in Figure 1(E, F) were done according to Joven et al., 2018 [70]: in brief, gelatin-embedding and colder temperatures (-30 °C) were used for sectioning.

For paraffin embedding, whole eyes were washed 3 times in PBS and fixed overnight in 10% formalin at 4°C. For intact lenses or later time points (100 dpl) when the lens is large, eyes were fixed in methanol: acetic acid (3:1 ratio) overnight at 4°C to preserve lens morphology. Paraffin embedded tissues were sectioned at 10μm thickness using a microtome.

### Immunofluorescent staining

Slides with cryo-sectioned eyes were air-dried for 30 minutes at RT and thereafter washed 3 times with PBST (0.2% Triton in 1 L PBS) for 10 minutes. To make the tissue more accessible to antibodies, a retrieval procedure was used. Plastic containers were filled with a citrate buffer (120 µL antigen unmasking solution (Vector, Peterborough, UK) in 11.88 mL PBS and preheated in a water bath to 86 °C. Slides were introduced once the solution reached 86°C and incubated for 10 minutes. Thereafter, slides were placed in a glass container with PBS at room temperature to cool down. Blocking buffer (10% goat or donkey serum in 0.2% PBST) was added to the slides for 1 hour at RT. Subsequently, primary antibodies were added to the slides and incubated overnight at 4 °C. List of primary antibodies and concentrations used can be found below. Slides were washed 3 times with PBST for 10 minutes. Appropriate secondary antibodies (AlexaFluor 488 or 594 conjugates, ThermoFisher) were added to the slides and incubated for 2-4 hours at RT protected from light. Slides were washed 3 times with PBST for 10 minutes. Hoechst (1:10000 in PBS) was added for 15 minutes at RT and protected from light. Slides were washed 3 times with PBST for 10 minutes and mounted with Fluorescent Mounting Medium (Sigma, #F-4680). Immunofluorescent staining procedures for the phagocytic experiments depicted in Figure 1(E, F) were done according to Joven et al., 2018 [70]. The same protocol was followed for paraffin embedded tissues, with additional deparaffinization steps. The deparaffination steps involved two xylene washes for 5 minutes, and gradually rehydration of the tissue by 1 minute washes with 100%,95%,80%70%,50%,30% ethanol followed by three PBS washes. All animals used for these experiments were post-metamorphotic, juveniles at 6-8 months old with an exception of the MPEG transgenic animals that were 3 years old.

<u>Antibodies:</u> ⍺-A-Crystallin (Gift by G. Eguchi, no dilution), Phospho-Histone H3 (Millipore-Sigma, #06-570, 1:200), eGFP (Abcam, #183734, 1:500), F4/80 (BioRad - Cl:A3-1 MCA497, 1:100), goat anti-GFP (Abcam, ab6673, 1:500), L-Plastin (LSBio, #LS-C344622, 1:100), P-CSF1R (Cell Signaling, #3154S,1:100), ⍺-SMA (Abcam, #5694, 1:100).

### Histology and Cytochemistry

Following deparaffinization, hematoxylin and eosin, and picrosirius red staining (Polysciences, #24901) were performed following manufacturer protocols. Prior to EdU and TUNEL assays, sections were deparaffinized, and incubated with 0.01M Sodium Citrate (pH=6) for 15 min at 95°C for antigen retrieval. A permeabilization step was followed in which the tissue sections were washed with 1% saponin and APBST wash buffer (APBS supplemented with 0.1% Triton-X100). Slides were then incubated in EdU (Invitrogen, #C10337) or TUNEL (Roche #11684795910 or #12156792910) reaction cocktails according to manufacturer guidelines. Slides were then washed 3 times in PBST for 10 min and nuclear counterstain was achieved by incubating slides with Hoechst 33342 (Invitrogen, #H3570) or DAPI (Life technologies, #D1306) at 1:1000 dilution. Slides were then washed in PBST 3 times and mounted with fluorescent mounting media (Sigma, #F-4680).

### Microscopy and imaging analysis

Live imaging of larvae, screenings and time-lapse movies were performed using either a Zeiss Axiovert 200 M inverted microscope or a LeicaM205 FCA stereo fluorescence microscope equipped with a Leica DMC 6200 camera. Figure 1 A, B images were produced by processing a Z-stack acquisition using the function ”Extended Depth of Focus” of LAS X version 3.7.4 software. Confocal images were obtained using Zeiss 700 and 710 Laser Scanning Confocal System Carl Zeiss, Gottingen, Germany). Z-stack configurations (Figure 1E: 26 images at 1µm intervals and 2048 x2048 size Figure 1F: 10 images at 1µm intervals and 1024x1024 size) were used to obtain high resolution images using ZEN 2012 Browser (Carl Zeiss, Gottingen, Germany). Fluorescent imaging was performed using a Zeiss Fluorescence Stereomicroscope Axio Zoom.V16 (Carl Zeiss, Oberkochen, Germany). ImageJ was used for image analysis.

### EdU+ cell quantification and statistical analysis

To determine whether the number of EdU+ cells in proportion to the total number of HOECHST+ cells give evidence of a difference between groups, we used Negative Binomial regression on EdU+ cell count with the two-level treatment as the predictor (PBS Liposome vs. Clodronate Liposome, Figure 3E; Clodronate/PBS bead vs. Clodronate/FGF bead, Figure 5B) and the number of HOECHST+ cells as the offset [139–141]. This model estimates the ratio of the mean EdU count to the total number of HOECHST+ cells for treatment versus control. Note that we used Negative Binomial regression because the measurement of interest is the number of EdU cells in proportion to the total number of HOECHST+ cells. These are counts rather than continuous numeric measurements, so an analysis based upon a count distribution is more appropriate than the more commonly used normal distribution-based procedures like the t-test. We used Negative Binomial regression rather than Poisson regression because there was evidence of overdispersion in the datasets. Because there were 4 eyes assigned to each treatment, we used a total of 8 EdU counts (and the associated HOECHST counts) to fit the model. Note that the Negative Binomial procedure is only approximate, since we have small sample sizes, and that by using the number of HOECHST+ cells as an offset, we are conditioning on their value. All animals used for these experiments were post metamorphotic, juveniles at 6-8 months old.

## Data Availability

Any code used in this study, as well as data produced, including transcript sequences and expression data, will be made available upon reasonable request to the authors.

## Resources availability

Both the plasmid Tol2-mpeg1:eGFP-polyA and the transgenic newt line *Pleurodeles waltl* tgTol2(Dre.*mpeg1:eGFP*)^MHY/SIMON^ are available to the scientific community upon request.

## Acknowledgments

The authors thank Laboratory Animal Resources (LAR) personnel and the Center of Advanced Microscopy and Imaging (CAMI). We acknowledge and thank the staff (Dr. Andor Kiss & Ms. Xiaoyun Deng) of the Center for Bioinformatics & Functional Genomics (CBFG) at Miami University for instrumentation and computational support. Special thanks to Myriam Aouadi for generously gifting us GeRPs to perform phagocytic assays. The authors further thank Dr. Jens Mueller and the staff from Research Computing Support for assistance with Miami’s Redhawk High Performance Computing cluster at Miami University. In addition, Lake Ernst for help on quantification analysis.

This research was supported by grants from the National Eye Institute: RO1 EY027801 and R21 EY033916 (to KDRT), R21 EY031865 (to HW and KDRT), by the John W. Steube Professorship Endowment (to KDRT), the Fight for Sight grant (to ASa), by the Miami University Doctoral Undergraduate Opportunity scholarship (to GT and SR), by National Institute of Neurological Disorders and Stroke grant F99 NS129167 (to JAT), by the Swedish Research Council and Cancerfonden (to ASi), by the Erasmus scholarship (to KJV), by Deutsche Forschungsgemeinschaft grants (DFG 22137416, 450807335 & 497658823) and TUD-CRTD core and seed funds (to MHY) and by a Karolinska Institutet Research Grant-Projektbidrag: FS-2020-0007 (to AJA).

