## Supplemental Material for "Macrophages modulate fibrosis during newt lens regeneration"

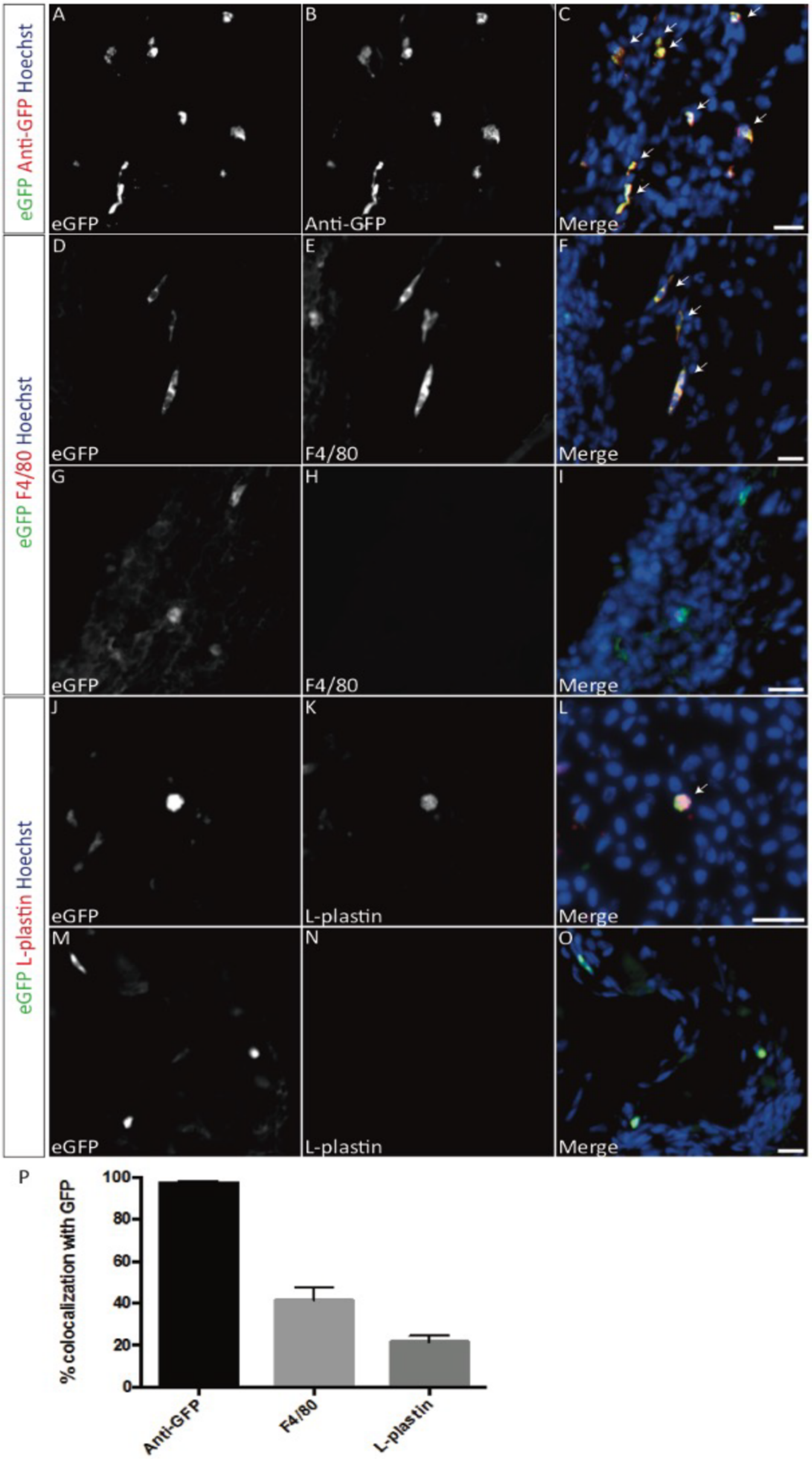
eGFP Anti-GFP Hoechst
eGFP F4/80 Hoechst
eGFP L-plastin Hoechst
A
eGFP
B
Anti-GFP
C
Merge
D
eGFP
E
F4/80
F
Merge
G
eGFP
H
F4/80
I
Merge
J
eGFP
K
L-plastin
L
Merge
M
eGFP
N
L-plastin
O
Merge
P
% colocalization with GFP
100
80
60
40
20
0
Anti-GFP
F4/80
L-plastin

**Supplementary Figure S1. *mpeg1:GFP* transgenic newts enable the *in vivo* labeling of macrophages.** (**A, D, G, J, M)** Representative fluorescence images of sections of 5-month-old *mpeg1:eGFP* newts showing presence of eGFP+ cells in the tail, trunk and head sections (**B,**) Anti-GFP immunofluorescence staining. (**C**) Merge of *mpeg1:eGFP* endogenous fluorescence, anti-GFP and Hoechst. (**E, H**) F4/80 immunofluorescence staining. (**F, I**) Merge of *mpeg1:eGFP*, F4/80 and Hoechst. (**K, N**) L-plastin immunofluorescence staining. (**L, O**) Merge of *mpeg1:eGFP*, L-plastin and Hoechst. Arrows represent colocalization events. (**P**) Percentage of colocalization of endogenous eGFP (average from tail, trunk and head sections) with anti-GFP (97.5%), F4/80 (41.4%) and L-plastin (21.6%). Scale bar: 50 µm; n=3. Related to Figure 1.

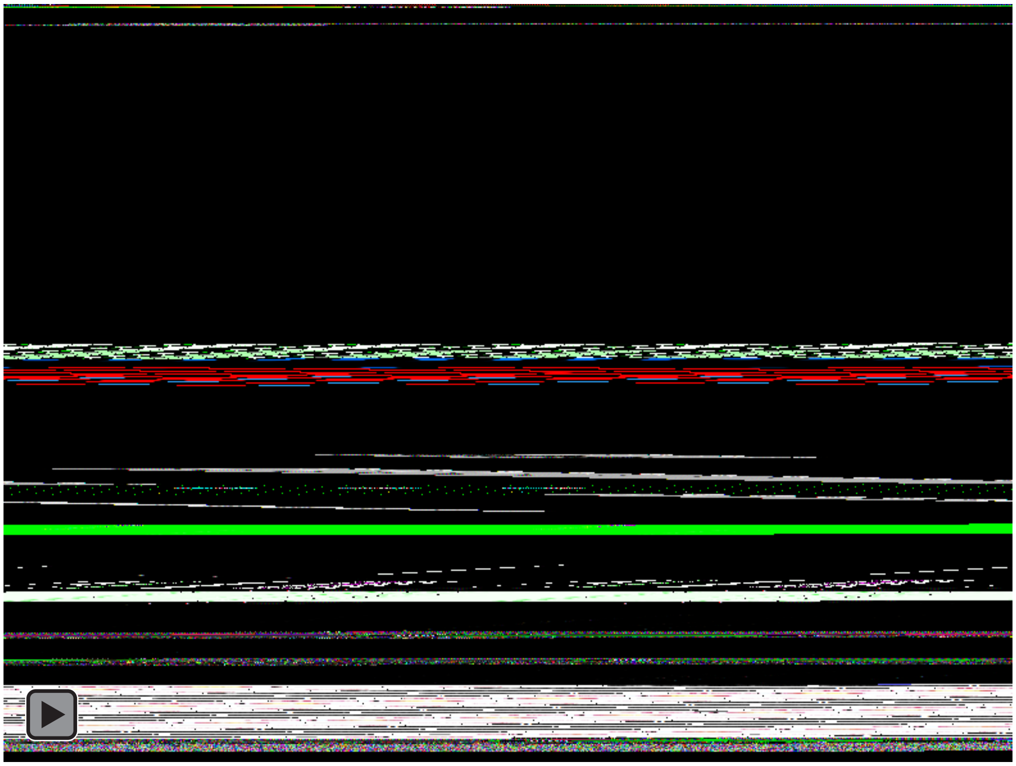

**Supplementary Movie 1. Time-lapse imaging of eGFP fluorescence in mpeg1:GFP transgenic newts.** Movie of the dorsal view of an $F_2$:mpeg1:GFP+ embryo. Positive macrophages and microglia of different morphologies are widespread. Border-associated macrophages can be seen floating in the cerebrospinal fluid inside the 4th ventricle. Related to Figure 1A, B.

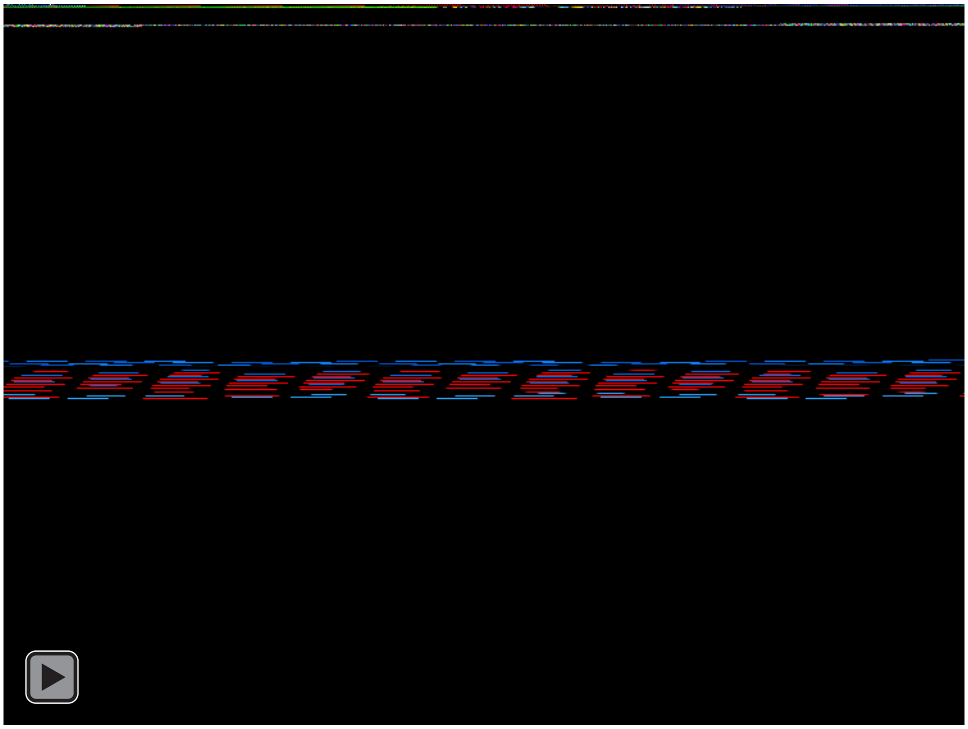

**Supplementary Movie 2. Phagocytic activity of eGFP + cells in the brain.** *In vivo* observation of GeRPs right after intraventricular injection in an $F_1$:mpeg1:GFP larva. Arrowhead point to the first events of GeRPs internalization by Mpeg1:GFP+ cells. Time-lapse images used to produce this movie were acquired during a total time of 5 minutes. Related to Figure 1D, E.

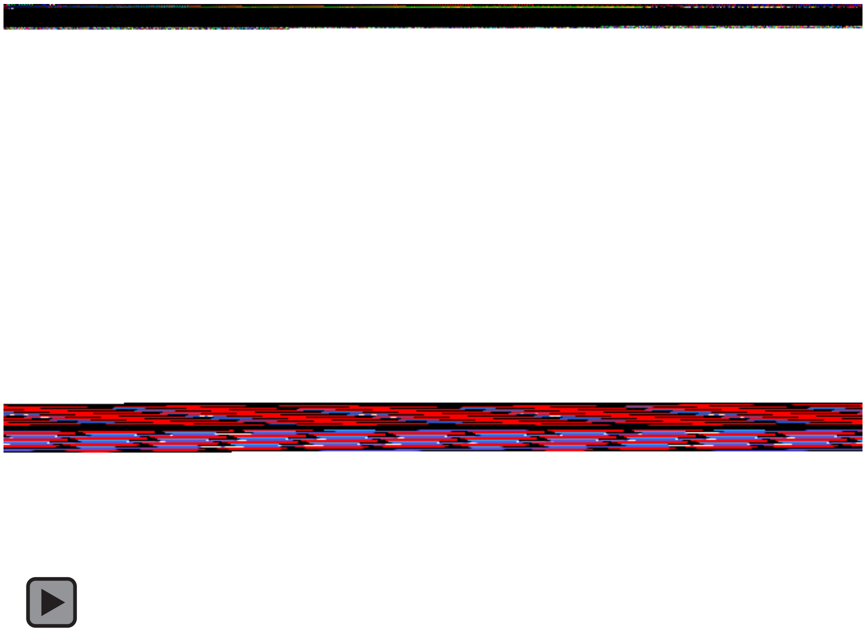

**Supplementary Movie 3. Phagocytic activity of eGFP + cells in the spinal cord.** *In vivo* observation of GeRPs after intraventricular injection in an $F_1$:mpeg1:GFP larva. Arrowhead point to an mpeg1:GFP+ cell approaching a single GeRP to potentially initiate phagocytosis. Time-lapse images used to produce this movie were acquired during a total time of 10 minutes. Related to Figure 1D, E.

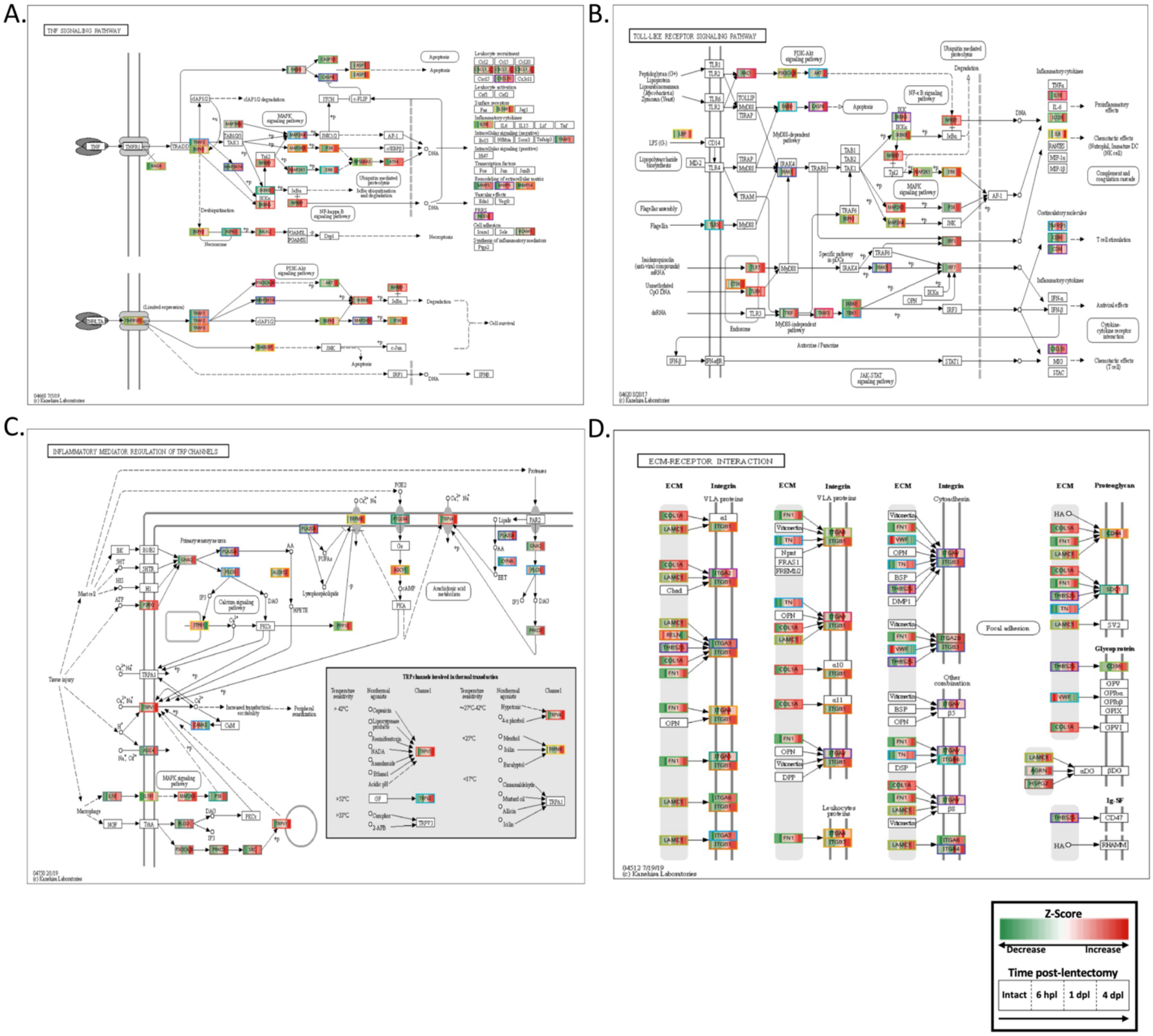


**Supplementary Figure S2. Time-dependent regulation of KEGG pathways in the injured dorsal iris of *Notophthalmus viridescens*.** A time course expression analysis was performed of the dorsal iris through 4 dpl. The shown pathways were overrepresented amongst transcripts that exhibited time-dependent regulatory patterns. The displayed expression values in each box represent the expression of homologous transcripts, ordered from left to right by time beginning with the intact iris. Color scale represents Z-score of expression values. Related to figure 2B.

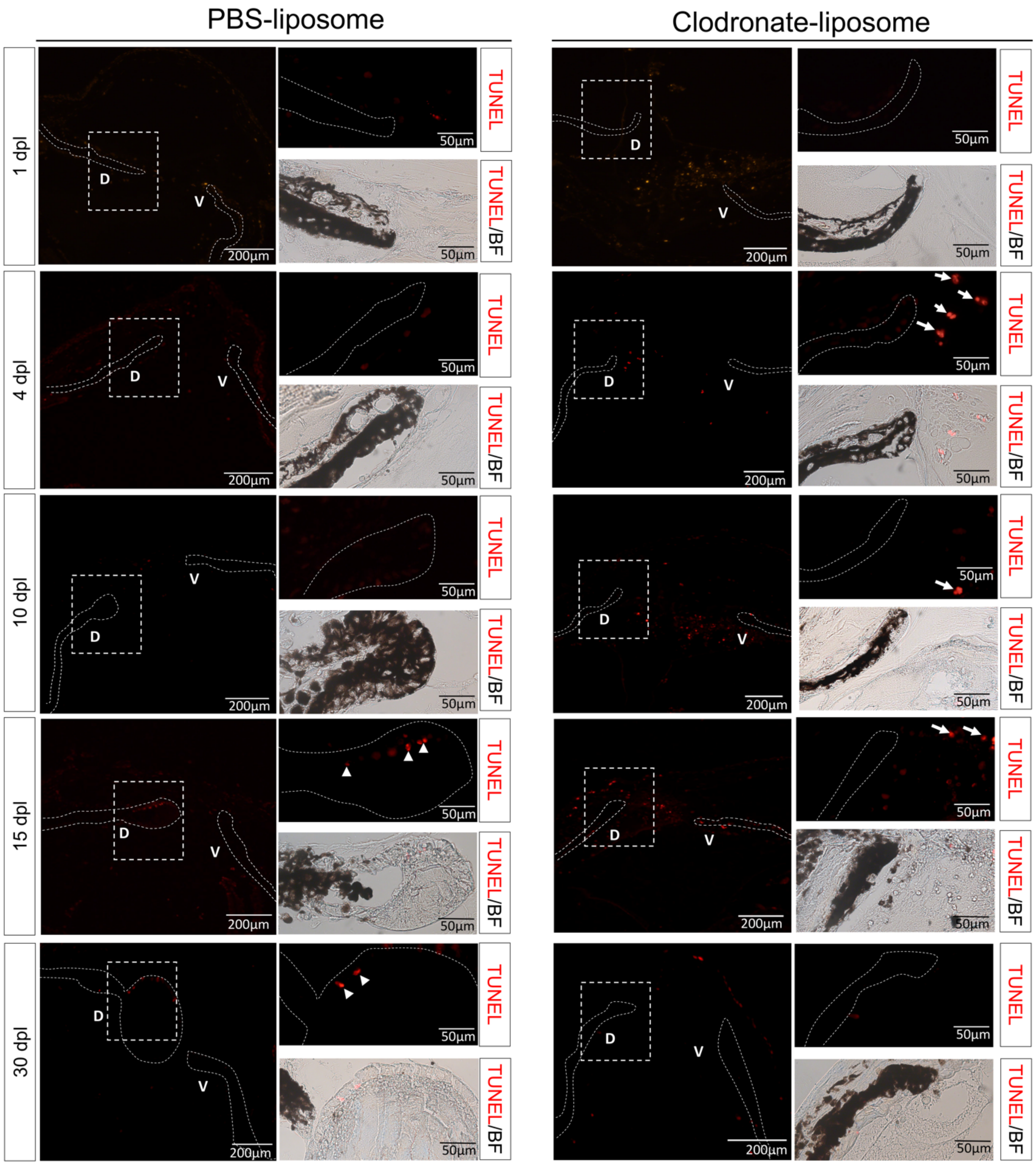


**Supplementary Figure S3. Clodronate treatment does not affect the survival of iPECs during the early stages of lens regeneration.** (A)TUNEL assay was used to visualize apoptotic nuclei from control- and clodronate-treated animals at 1, 4, 10, 15, and 30 dpl (paraffin embedded tissue). Dashed lines were used to mark the iris epithelium. Inset images of the dorsal iPECs highlight the effects of macrophage depletion on cell survival. As expected, TUNEL+ nuclei were observed in the vitreous and aqueous chambers of clodronate-liposome treated eyes (arrows) but not in PBS-liposome treated eyes. At 15 (Stage IV-V) and 30 dpl (Stage VIII) TUNEL+ nuclei were found in the lens epithelial layer of PBS-liposome treated eyes (arrowhead); n=6 per time point. Scale bars: 200µm (overviews, left) and 50µm (insets, right).

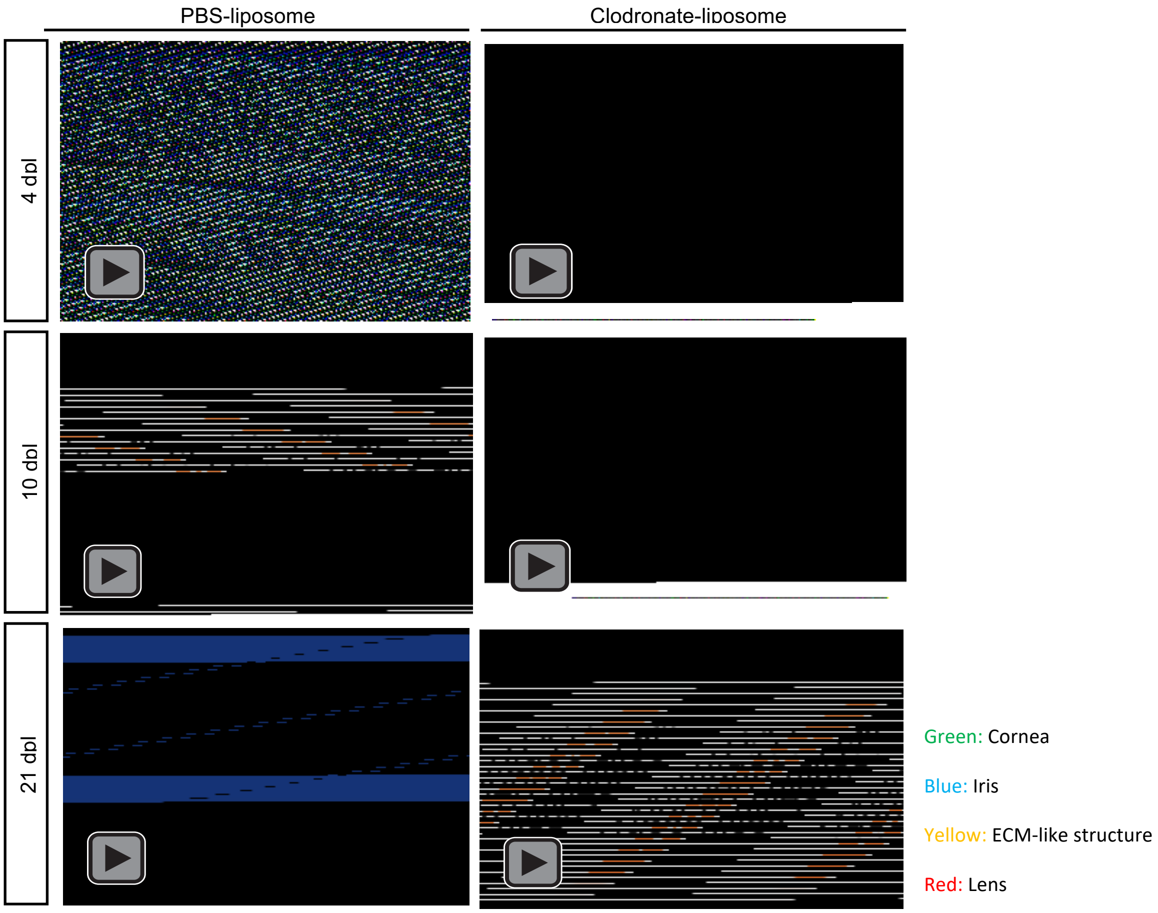


**Supplementary Figure S4. Three-dimensional representation of OCT images from PBS and clodronate treated eyes.** Animated rendering of three-dimensional images, reconstructed from OCT C-scans. Eye tissues (cornea/green, iris/blue, ECM/yellow, regenerating lens/red) were manually pseudo colored to aid in visualization. In PBS-treated eyes, ECM accumulation is observed at 4 dpl (Stage 0-I) and cleared by the 10 dpl (Stage I-II) prior to the formation of the new lens vesicle at 21 dpl (Stage VI-VII). ECM in clodronate treated eyes was more abundant at 4 dpl and failed to resolve by 21 dpl; n=10.

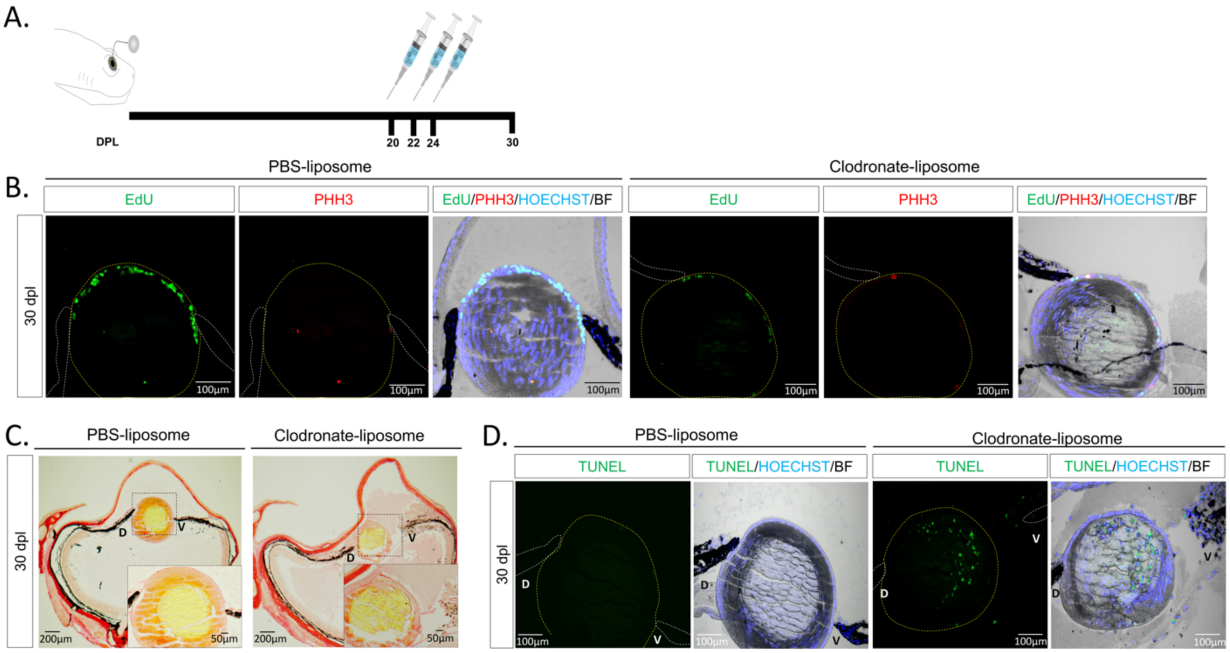


**Supplementary Figure S5. Late clodronate-liposome administration impairs lens growth by increasing apoptosis instead of affecting proliferation. (A)** Schematic representation of experimental design. Clodronate or PBS liposomes were injected intraocularly in the aqueous chamber at 20, 22, and 24 dpl in the presence of the regenerating lens. **(B)** Clodronate liposome administration at 20 dpl did not inhibit the proliferation and mitosis levels of lens epithelial cells; n=6. Scale bars: 100µm (paraffin embedded tissue). **(C)** Picrosirius red staining revealed a stronger collagen staining (red) in the vitreous and aqueous chamber of the clodronate liposome treated eyes; n=6. Scale bars: 100µm (paraffin embedded tissue). **(D)** Apoptotic cells were detected inside the lens fibers and at the surrounding area of the ventral iris following macrophage depletion at 20 dpl; n=6. Scale bars: 100µm. Related to figure 7 (paraffin embedded tissue).

| Gene | Forward Primer | Reverse Primer | PCR product (bp) | Target Sequence |
|---|---|---|---|---|
| P53 | TATGGCACCACCACGCTATG | AATGATGGTCATGCTCCCCC | 107 | M1034543_PLEWA04 |
| CDK2 | CGGTATCCCTTTGCCACTCA | TCAGCAAGCTTGATAGCCCC | 144 | M0441300_PLEWA04 |
| E2F1 | TGCCGGCCAAAAGAAAGTTG | CTATTCCGGTCCGGCCTTTT | 90 | M0222145_PLEWA04 |
| SOCS3 | TACATGCCAAACAGCGGACT | GTGCCCGTTGACAGTTTTCC | 162 | M0047129_PLEWA02 |
| CSFR | TGGCACTGATAGTCGCAGTC | AAACACCGGTTCCCCTTTGT | 101 | M0373017_PLEWA04 |
| COX-2 | GGGAGCTTTGATTTTCGCCC | CGGTCATACACACTCCTCGG | 92 | M0026016_PLEWA04 |
| IL1b | CCGCAGGATAGTGGTGATCG | TGTCCCCAAACAGACACCTG | 77 | M0346665_PLEWA04 |
| TGFB2 | AAACCCAGAAGCATCAGCCT | GCTGCACTTGCAGGACTTTAC | 133 | M0018968_PLEWA04 |
| TGFB3 | AACGCTTCATCAGTGGGAGG | CACTCCCGCACATTCTCTGT | 85 | M0100785_PLEWA04 |
| MMP3/10a | CAGGCTGAGAGGGAGAGTCA | GCGCACACTAAGAGGAGTGA | 102 | M1347552_PLEWA04 |
| MMP9 | GAGGGGGTGCAGTACCTCTA | AGCTGTTACTGGGGTTGTCG | 111 | M0114547_PLEWA04 |
| TAII70 | CCTGGGAAGCATTTGGTAGA | TTCACGAGCTGTCTGTGGAG | 137 | M0451743_PLEWA04 |

Table S1. Oligos and target sequences used for RT-qPCR
